# A methodology for morphological feature extraction and unsupervised cell classification

**DOI:** 10.1101/623793

**Authors:** Dhananjay Bhaskar, Darrick Lee, Hildur Knútsdóttir, Cindy Tan, MoHan Zhang, Pamela Dean, Calvin Roskelley, Leah Edelstein-Keshet

**Affiliations:** Department of Mathematics, University of British Columbia, Vancouver, BC, Canada; Department of Cellular & Physiological Sciences, University of British Columbia, Vancouver, BC, Canada

## Abstract

Cell morphology is an important indicator of cell state, function, stage of development, and fate in both normal and pathological conditions. Cell shape is among key indicators used by pathologists to identify abnormalities or malignancies. With rapid advancements in the speed and amount of biological data acquisition, including images and movies of cells, computer-assisted identification and analysis of images becomes essential. Here, we report on techniques for recognition of cells in microscopic images and automated cell shape classification. We illustrate how our unsupervised machine-learning-based approach can be used to classify distinct cell shapes from a large number of microscopic images.

**Technical Abstract:** We develop a methodology to segment cells from microscopy images and compute quantitative descriptors that characterize their morphology. Using unsupervised techniques for dimensionality reduction and density-based clustering, we perform label-free cell shape classification. Cells are identified with minimal user input using mathematical morphology and region-growing segmentation methods. Physical quantities describing cell shape and size (including area, perimeter, Feret diameters, etc.) are computed along with other features including shape factors and Hu’s image moments.

Correlated features are combined to obtain a low-dimensional (2-D or 3-D) embedding of data points corresponding to individual segmented cell shapes. Finally, a hierarchical density-based clustering algorithm (HDBSCAN) is used to classify cells. We compare cell classification results obtained from different combinations of features to identify a feature set that delivers optimum classification performance for our test data consisting of phase-contrast microscopy images of a pancreatic-cancer cell line, MIA PaCa-2.

## Introduction

The morphology of cells often reflects their tissue-specific function or state. Cells undergo morphological changes to become motile in response to various stimuli, either as part of normal physiology and development, or due to pathologic disorder. In some cancers, epithelial cells lose their inter-cellular connections, and take on protrusive, exploratory morphologies as they transit from static to migratory phenotypes. Recognizing such abnormal cell shapes can aide in properly identifying malignant (as distinct from benign) tumors. Yet, this analysis of cell shapes derived from microscopic images has traditionally been handled by expert humans, i.e. pathologists and histologists, who use many cellular features, including cell shapes, in their diagnoses. Modern medicine and high throughput quantitative biology have motivated new methods for extracting useful information from microscopic data [1]. In particular, data science and machine learning is being applied to devise new intelligent computational tools to process microscopic images and recommend patient-specific clinical treatments.

Two broad classes of machine learning are the supervised and unsupervised methods. Both can be used to address classification and regression problems. Supervised learning algorithms require training data (annotated subset of data for which the desired result or ground truth is known *a priori*) to train a classifier or regressor. Typically, the learned classifier or regressor retains sufficient generality to handle new data for which the outcome (category in the case of classification) is unknown. Several examples of this type include the following. Keskin et al [2] used dual-tree complex wavelet transform to classify cell images into 14 types of known breast and liver cancer types (i.e. match images to predetermined classes). Similarly, Reta et al [3] developed automated methods to classify leukemia cell images to one of four leukemia types based on morphology of their nuclei and cytoplasm. Matsuoka et al [4] used morphological features of stem cell microscopy images and training sets to predict the differentiation stat eof the cells, and verified the results using biochemical markers. (Here we eschew training data.) Similarly, Nanni et al [5] classified stem cells into three stages of maturation using texture descriptors using wavelets, Gaussian filters, and other methods. Park et al [6] used automatic processing of microscopy images to compare morphologies of prostate cancer cells from patients and from cell cultures. Many of these papers share some features of segmentation or clustering methods with our techniques, but with an important distinction: In these examples, there exists a priori knowledge of the basic classes of objects to be identified i.e. the goal is to match cell features with known categories. Our goal is to extend such methods to datasets where no such pre-knowledge is available.

Unsupervised learning algorithms exploit patterns inherent in the data to make predictions under the premise that similar data should correspond to similar outcomes. In practical applications, training data is often unavailable and requires significant time, expense and effort to acquire. This is a limitation that can be avoided by using unsupervised learning methods, and motivates our approach in this paper.

Here we develop a software-based pipeline for delineating cells boundaries in micrographs (a process known as microscopic image segmentation), quantification of cell morphology (called feature extraction in machine-learning parlance) and subsequent cell shape classification. The pipeline operates in a semi-supervised manner, with user intervention required only at the segmentation stage to pick out correctly segmented cells. Alternatively, user assistance is required to train a supervised algorithm to perform image segmentation. The remainder of the pipeline (for feature extraction and classification) is automated and does not require any user input.

We illustrate our methodology using phase-contrast images of pancreatic carcinoma cells (MIA PaCa-2 cell line), though the methods are general and suitable for other types of cell images. MIA PaCa-2 is a human cell line that was established by A. Yunis, et al. [7] from primary tumor tissue of the pancreas. It is currently used as an *in-vitro* model to study carcinogenesis in pancreatic ductal adenocarcinoma [8]. MIA PaCa-2 cells exhibit several distinct morphological shapes, making them particularly suitable for our purposes. Since our methodology does not require biomarker cell labeling, it can be easily adapted for confocal or fluorescent images. Phase microscopy is chosen for demonstration purposes because phase images are particularly challenging to segment [9].

We assemble a pipeline using unsupervised learning methods that uses inherent properties of the input data to extract clusters and determine a natural classification. Each cell is associated with a feature vector, whose elements (“features” or “descriptors”) quantify aspects of its morphology. Feature selection varies with imaging modality (bright-field, phase-contrast, florescent, etc.) and target application. Therefore, we compute a diverse range of features and evaluate their performance on the MIA PaCa-2 data set to decide which features provide the best result.

Unsupervised methods are technically challenging to design and implement, especially in the context of image classification, but have the advantage that they are self-contained and require no expert curated training data. Unlike state-of-the-art supervised methods (e.g., convolutional neural networks [10]) where feature computation is an inherent part of training, unsupervised methods can only take advantage of a set of pre-determined features. According to a recent review, supervised methods outperform unsupervised clustering methods despite the availability of large amounts of data, due to lack of consensus in class delineation, sensitivity to algorithm parameters, limitations in feature extraction and lack of robustness in cluster identification [11]. However, by computing a large number of features, performing automated feature selection and limiting the number of output categories, this shortcoming can be partly overcome. Additionally, features computed as part of the unsupervised classification methodology contain information that can be easily interpreted by biologists and medical professionals.

Despite their challenges, unsupervised learning methods have been gaining popularity. Logan et al. [12] devised a learning methodology based on pixel intensities of fluorescent cell images as features. Gençtav et al. [13] used pixel brightness along with measurements of area, perimeter, shape factors and Feret diameters for both nuclear and cytoplasmic segmentations to identify normal and malignant cervical cells. Segmentation was performed in a semi-supervised manner where a single boundary could be shared by more than one cell in cases where cells overlapped. Ahonen et al. [14] used unsupervised clustering methods to classify simulated tumor and prostate cancer spheroids. They used geometrical features calculated from ellipse fitting, boundary features obtained from principal curve fitting and texture-based features.

Kriegel et al. [15] classified the 3-D surface of myeloid cells by computing Fourier shape descriptors from a series of 2-D projections and by using a Self-Organizing Map (SOM, an unsupervised competitive learning method based on artificial neural networks) to perform classification in the Fourier domain. Alizadeh et al. [16] analyzed shapes of osteosarcoma cells in pairs of cell lines with distinct invasion potential. They used Zernike moments as feature vectors. Their method succeeded in assigning cells into their respective invasiveness categories. As in Fourier analysis, the original image can be fully reconstructed by superposition of an arbitrarily large collection of its Zernike moments. However, the individual moments do not, themselves, provide intuitive descriptors that biologists can easily appreciate.

In this paper, we only consider morphological features and exclude texture information. This enables us to develop methods that are more generic and do not depend heavily on the type of microscopy image. Furthermore, we restrict our approach to 2-D images as proof of principle. In principle, the same approach can be used to classify segmented 3-D cell shapes, where the 3-D segmentation is obtained from a volumetric image (acquired using a confocal micropscope for instance) using existing methods such as the marching cubes algorithm. Our goal is to develop unsupervised classification of cells based solely on morphological features, requiring no additional cell labeling or staining. While this makes for a more challenging classification process, it accommodates a far wider domain of acceptable input data, allowing for live-cell imaging without artifacts of fixation, staining or fluorescent labeling.

Our methodology, while illustrated on MIA PaCa-2 cells, is applicable to a wide range of cells and image types. Many related label-free methods currently used do not achieve multi-class classification [17]. Conversely, other methods cannot process high-throughput data due to computational complexity of feature extraction. Therefore, a major objective of our work is to select from a diverse pool of easy to compute features that are capable of classifying cells based on their morphology.

## Methods

In this section, we describe our methodology for unsupervised classification of Mia-PaCa2 cells (American Type Culture Collection, Manassas, VA, Cat # CRL-1420) maintained in sub-confluent 2-D monolayer culture and imaged using phase contrast microscopy (Nikon, TMS microscope using a 10X objective). Typically, the machine learning algorithm is embedded into a processing pipeline that converts microscopy images into numerical data corresponding to individual cells [18]. The pipeline consists of image processing, feature extraction, dimensionality reduction, and classification steps, as described in detail below. Due to the absence of labels in our data, validation of our results is performed by human experts (Pamela Dean, Calvin Roskelley, Dhananjay Bhaskar) who judge the performance of each classifier.

The first phase of image processing is to enable separation of foreground and background by removing artifacts, reducing noise and compensating for uneven illumination. Subsequently, image segmentation methods (techniques that divide an image into regions of interest) are used to identify cells amongst the foreground pixels. The choice of segmentation algorithm depends on the image and cell type. No single algorithm is capable of identifying cells from diverse imaging modalities. Currently, our segmentation is semi-supervised, with a human expert selecting correctly segmented cells amongst the results of the image processing software.

For each segmented cell, a feature vector consisting of quantifiable descriptors of cell shape and size is computed. Each feature in the vector is normalized by subtracting its mean and dividing by its standard deviation, where these statistics are calculated over all segmented cells in the entire data set. Standardization of features in this manner (called Z-score standardization) is a frequently used method for comparing features on a common standard irrespective of the underlying distribution of feature values. A classification algorithm uses the normalized feature vectors to define and to distinguish between “cell types”. The performance of the classifier depends on the quality of segmentation and accuracy of features. Two cells can be distinguished (i.e. assigned different labels by the classifier) if their feature vectors are sufficiently different. Furthermore, in order to identify a specific morphology, one or more features must capture unique characteristics of that morphology.

Most widely used classification algorithms based on a distance metric between features perform well for low dimensional feature vectors. Dimensionality reduction techniques convert high dimensional feature vectors to lower dimensions by combining multiple features to remove redundancy while preserving variance in data. Two popular dimensionality reduction techniques, Principal Component Analysis (PCA) and t-distributed Stochastic Neighborhood Embedding (t-SNE) are described in the dimensionality reduction section below. After dimensionality reduction, clustering algorithms are used to classify data points corresponding to individual cells. Several clustering algorithms are publicly available and have been empirically evaluated using synthetic data for their performance, robustness and accuracy [19]. We use HDBSCAN (hierarchichal density-based spatial clustering of applications with noise) to identify clusters and quantify similarity between cells. We evaluate our classification results using different sets of features and dimensionality reduction techniques.

Figure 1 indicates steps in a standard data processing pipeline for object classification, which we have automated and adapted for cell shape classification. Our pipeline is modular, so that individual components can be changed or replaced as needed to accomodate different imaging modalities (fluorescent, brightfield, phase-contrast, multi-photon, DIC, etc.) and sample preparation methods.

**Figure 1.**
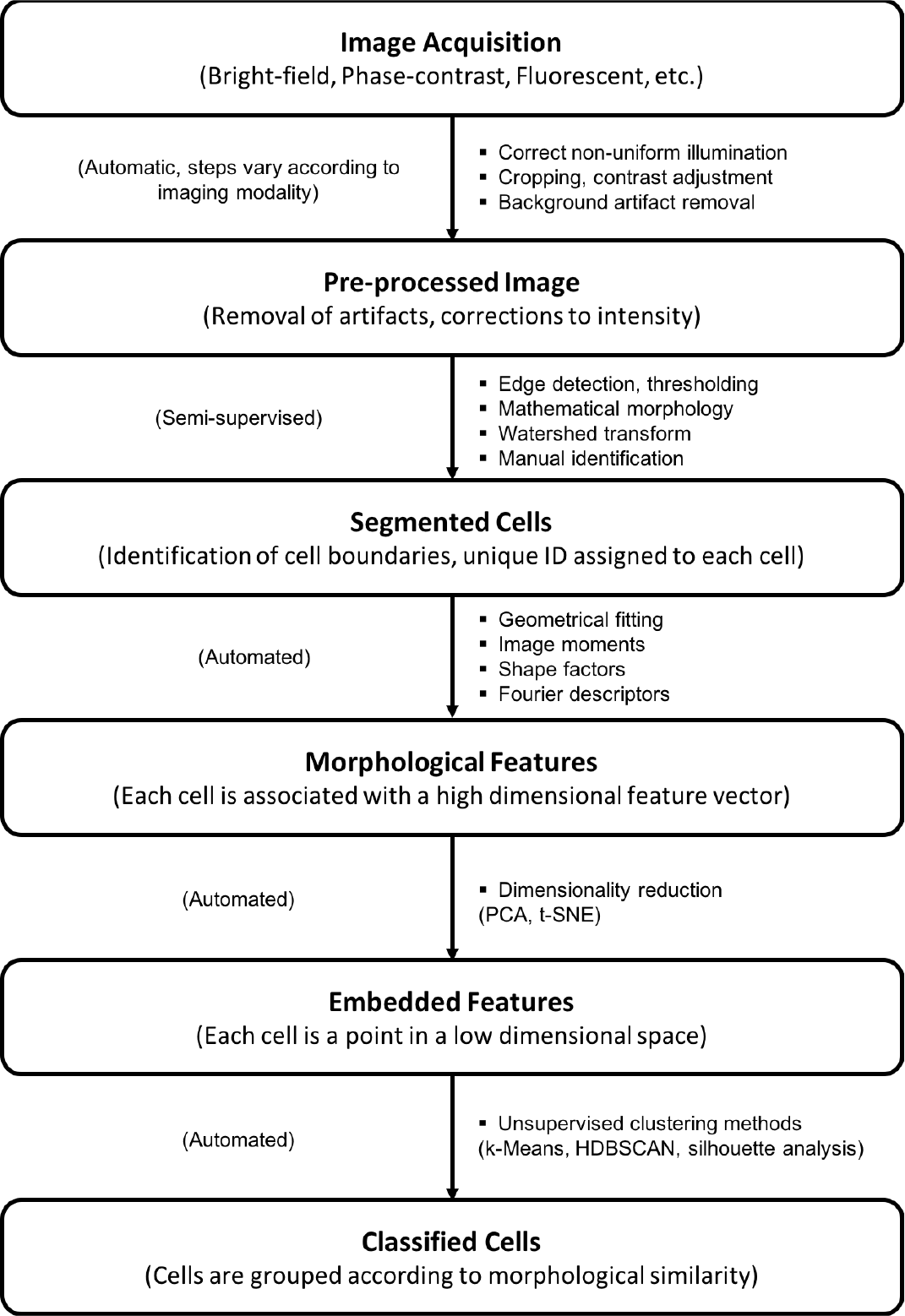
Standard pipeline for cell classification using microscopy images. The output from each stage is printed in bold. The bulleted lists are commonly cited methods used in the computation. Our methodology is predominantly automated, as indicated in parentheses on the left.

### Image Processing

Identifying individual cells in an image is essential for automated recognition of multiple cell types in large cell populations. Automated processing of 2-D images to count cells and identify cell types using morphological measurements has been steadily gaining traction since the 1960s. Over the past decades, literature on the subject has grown exponentially, according to a review published in 2012 [20].

Most current segmentation methods are based on basic approaches such as intensity thresholding, feature detection, morphological filtering, region accumulation and deformable model fitting (reviewed in [20]). We overcome issues of inaccurate cell boundaries and over-segmentation (typical in Voronoi-based or watershed transform methods) by bootstrapping watershed segmentation with markers lying inside cell boundaries. Popular deformable model approaches such as geodesic active contours or level sets to detect cell boundaries (by minimizing a predefined energy functional), can result in poor boundary detection: their local optimization algorithms can get trapped in local minima [21].

Segmentation of bright field and phase contrast images is generally more challenging compared to fluorescent images. The latter usually have better contrast and deformable model fitting techniques like active contour or level sets work well [9]. Distinctive bright white patches or halo surrounding cells in bright field and phase contrast images prevent accurate determination of cell boundary. Therefore, a custom approach is required for each application that takes heterogeneity in cell shape, population density, variability in cell compartmentalization, etc., into account.

The following sections describe our approach for segmenting phase contrast images using a combination of edge detection, thresholding, mathematical morphology and watershed transform.

#### Foreground Detection

Foreground detection is performed in three stages. First, the Sobel-Feldman derivative filter is applied to the original grayscale image to find edge points. These points are pixel locations in the image corresponding to non-zero intensity changes. The Sobel-Feldman operator uses two 3 × 3 kernels, one for derivative in a horizontal direction and the other for derivative in a vertical direction, which are convolved with the original image to calculate gradient approximation. The result is binarized by thresholding, with the value of the threshold specified by the user.

The binary image produced by edge detection is further manipulated using mathematical morphology, as shown in Figure 2. Mathematical morphology is a collection of set-theoretic operations on binary images that have been used for image enhancement, noise removal, edge detection, etc. Foundations of mathematical morphology are based on two operations: erosion and dilation. (See S1 Appendix for details.)

In the second stage of foreground detection, edge points are connected by dilating the image with line shaped structural elements. The size of the structural elements is a parameter set by the user. This leads to the formation of closed loops around isolated cells or clusters of tightly packed cells. The final stage involves filling of closed loops and removal of small objects and artifacts whose size is below the user-specified threshold for minimum cell size. Cells that cross the boundary of the image are also removed during this stage of image processing.

**Figure 2.**
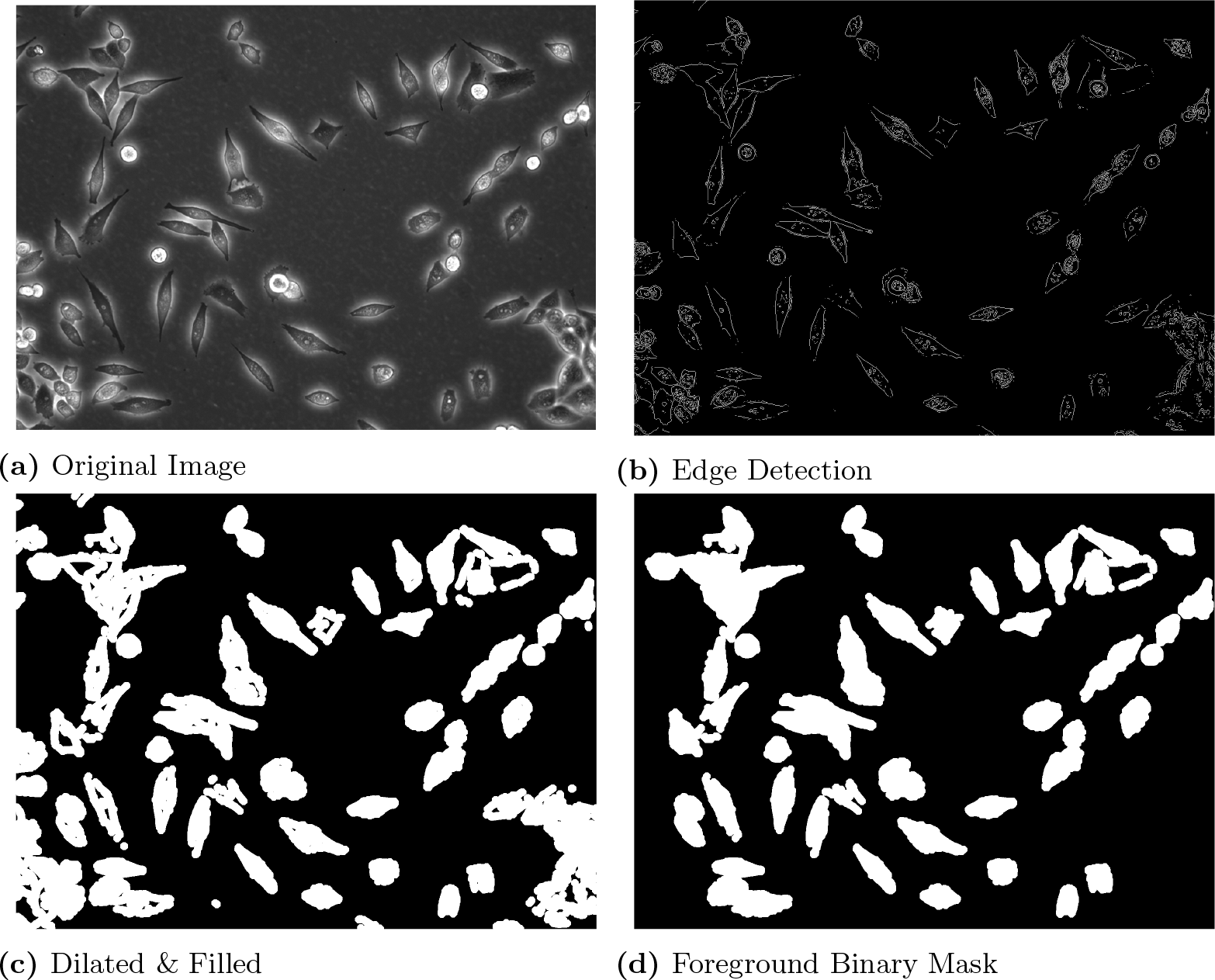
Foreground detection from phase contrast image. (a) Original phase contrast microscopy image of MIA PaCa-2 pancreatic carcinoma cells. (b) Edge point detection after application of the Sobel-Feldman derivative filter and conversion from grayscale to binary by thresholding. (c) Dilation by line-shaped structural element. (d) Resulting foreground markers obtained after filling, removal of small artifacts and objects connected with image boundary.

#### Cell Segmentation

Foreground detection separates foreground from background in the original image. The foreground binary image is eroded *N* times (with *N* and size of structural element specified by the user) to obtain foreground markers, a majority of which lie inside cell boundaries. The marker-based watershed transform is a region accumulation approach that segments cell boundaries using foreground markers and gradient of the original image. The number of correct segmentations in the result depends on the pre-processing of markers prior to the segmentation [22].

The watershed segmentation algorithm requires foreground markers (to identify regions inside individual cell) and background markers (for regions between adjacent cells). Over-segmentation may occur if background markers are too close to cell edges. To prevent this, we computed background markers using the “skeleton by influence zones” (SKIZ) of the foreground markers [23]. Pixels in the immediate vicinity of a given foreground marker (closer to it than to any other foreground marker) form its influence zone. SKIZ, the boundary between influence zones of all foreground markers, is analogous to the Voronoi tessellation of foreground markers in the image plane. In practice, background markers are determined by (1) computing the distance transform of the foreground markers and (2) finding ridge-lines using the watershed transform of the distance-transformed foreground markers (see Figure 3b); ridge-lines correspond to background markers or SKIZ. Given foreground and background markers, the priority-flood watershed algorithm is applied to the original image, resulting in watershed lines. These lines identify cell boundaries, see Figure 3c.

**Figure 3.**
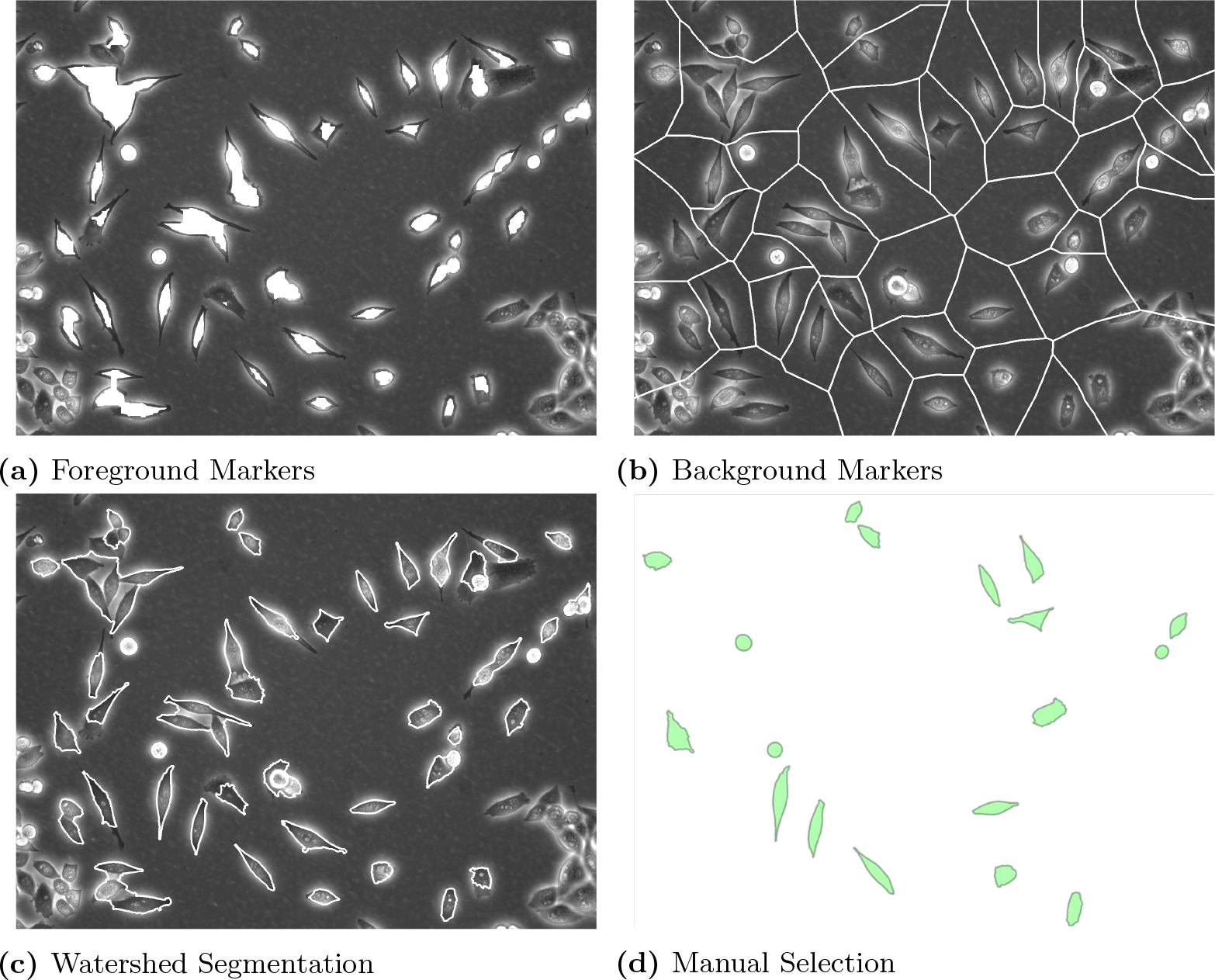
Cell segmentation. (a) Eroded binary foreground markers overlaid on top of original image. (b) Background markers computed by applying watershed transform to the distance transform of fore-ground markers. (c) Result of watershed segmentation using foreground and background markers. (d) Manual selection of correctly segmented cells.

The entire image segmentation process requires minimal user input to (1) specify the threshold parameters, size of structural elements and number of iterations for morphological operations and (2) to manually select correct segmentation after the watershed transform is applied, ensuring that only correctly segmented cells are assigned unique ID numbers (and serialized) for further processing in the pipeline.

### Feature Extraction

Both boundary and area-based methods are used to extract morphological features from segmented cell shapes. Both types of methods use cell-boundary data, but area-based methods also use cell-interior points. Area based methods are more robust to small perturbations in cell shape and are easy to compute. For example, to accurately estimate the area of a given shape it suffices to count pixels in its interior, whereas perimeter estimation is not so straightforward [24]. The main advantage of boundary based features (curvature functions, cubic spline interpolation of cell boundary, normalized Fourier shape descriptors, etc.), is that they provide a good quantization of angles, corners and curves in the image. Such geometric details are lost in many area-based features.

#### Hu’s Moment Invariants

Moments of a distribution are integrals that characterize means, variances, and higher order properties of density distributions such as distributed mass *m*(*x, y*), probability density *p*(*x, y*), or geometric shapes. In the case of binary cell images (e.g., Figure 2d), the distribution of interest is a function *f* (*x, y*) that takes points (*x, y*) in the plane into binary values, 0 or 1 (for points outside or inside the cell, respectively). The *pq*’th image moment (of order *p* + *q*), denoted *m*_*pq*_, is defined as:

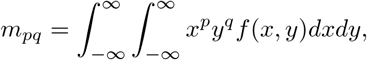

which are approximated, here, by the discrete sum over all pixels in the raster image:

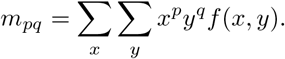

The zeroth raw moment, *m*_00_ is then the cell area, whereas 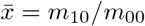 and 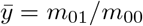 are coordinates of the cell centroid. For translational invariants, moments are typically computed relative to that centroid,

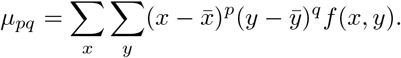

The zeroth central moment, *μ*_00_, is equivalent to *m*_00_ and corresponds to the area of a segmented cell.

In classifying cell shapes, it is desirable to assign shape-features that are invariant to image size, to rotation, and/or to reflection (mirror image). This can be accomplished by normalizing some moments, and by deriving others that are invariant to such operations. When the axes of a 2-D image are scaled by a factor *λ* (e.g., with *λ* > 1 for magnification, *λ* < 1 for reduction), the moments *μ*_*pq*_ of the unscaled image are transformed to

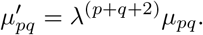

Hence, a normalized (scale-invariant) central moments *η*_*pq*_ is

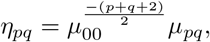

(obtained by setting 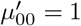, which scales cell area to unity.)

A set of rotationally-invariant moments *φ*_*i*_, *i* = 1 … 7 derived by Hu [25], are widely used for translation, scaling and rotation invariant pattern recognition, including recognition of typed English language characters. We adopt these moments (summarized in S2 Appendix) to describe cell shape features. One of the Hu moments, *φ*_7_, is skew invariant in addition to translation, scaling and rotational invariance. Unlike raw or central moments, *φ*_1_ … *φ*_7_ do not form a complete set of image descriptors. While higher order moments can be calculated, image reconstruction given a set of Hu’s moment invariants is not straightforward. Furthermore, all seven invariant moments are zero for images that are rotationally symmetric [26].

Dunn and Brown used shape measures (extension, dispersion and elongation) and principal axis orientation calculated using *φ*_1_ and *φ*_2_ to characterize the shape and alignment adopted by chick heart fibroblasts on micro-fabricated grooved substrata [27], but adoption of Hu’s moments has declined in recent literature on morphology-based cell classification. The role of these invariants and their usefulness is investigated in the results section.

#### Geometrical and Boundary Features

For high-throughput cell classification, the cell-shape feature vector should be concise and computationally inexpensive. While Hu’s moments meet this criterion, they are not easily interpretable in terms of intuitive features. Normalized Fourier shape descriptors (NFSDs) represent the boundary of an object using a subset of Fourier mode coefficients [28]. One benefit of computing NFSDs is that synthetic cell images can easily be generated by sampling from Fourier coefficients and computing the inverse transform. In cases with limited number of segmented cell images, these synthetically generated cell boundaries can be used to augment the origin data for classifier training. However, the number of coefficients required for reasonable accuracy is hard to estimate, is often large, and depends on the curvature of the object. Recent work in the Marée-Grieneisen lab [29] has led to a generalization of cell shape Fourier analysis called Lobe Contribution Elliptic Fourier Analysis (LOCO-EFA). This method has been used to quantify the shape of pavement-cells on plant leaves. Individual modes of LOCO-EFA are more directly interpretable in terms of actual cell shapes.

Point-wise values of curvature on the perimeter of a cell provide a detailed description of its shape. For example, peaks in the curvature correspond to corners or tips of thin protrusions. Urdiales et al. [30] describe a non-parametric method for efficient computation of a curvature function for an object contour, represented as a short feature vector. Their method requires pre-computation of certain Fourier transforms for typical shapes, which is an obstacle to high-throughput cell classification (particularly where the entire data set is not available in advance).

In view of the above, we bypass the use of curvature functions. Instead, we use simple fitting of conics (circles and ellipses), which are universally applicable to all planar objects. A set of boundary points (obtained from segmentation) is used as input and uncertainty is quantified from goodness of fit measures. Cell boundary descriptors are obtained from cubic spline interpolation, with the number of spline points estimated using a manually adjusted smoothing parameter. Together, these descriptors encode information about the shape and size of the cell that is easy to visualize (Figure 4). Like Hu’s moments, these features are invariant to rotation and translation, as well as to noise in the shape boundary.

Consider an arbitrary curve *f*(*θ*) = 0 parametrized by *M* features, *θ* = (*θ*_1_, *…, θ*_*M*_)^*T*^. We fit this geometry to a set of boundary points 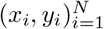, bysolving the following optimization problem:

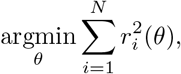

where *r*_*i*_ is the orthogonal distance between boundary point (*x*_*i*_, *y*_*i*_) and the shape *f* (*θ*) = 0.

**Figure 4.**
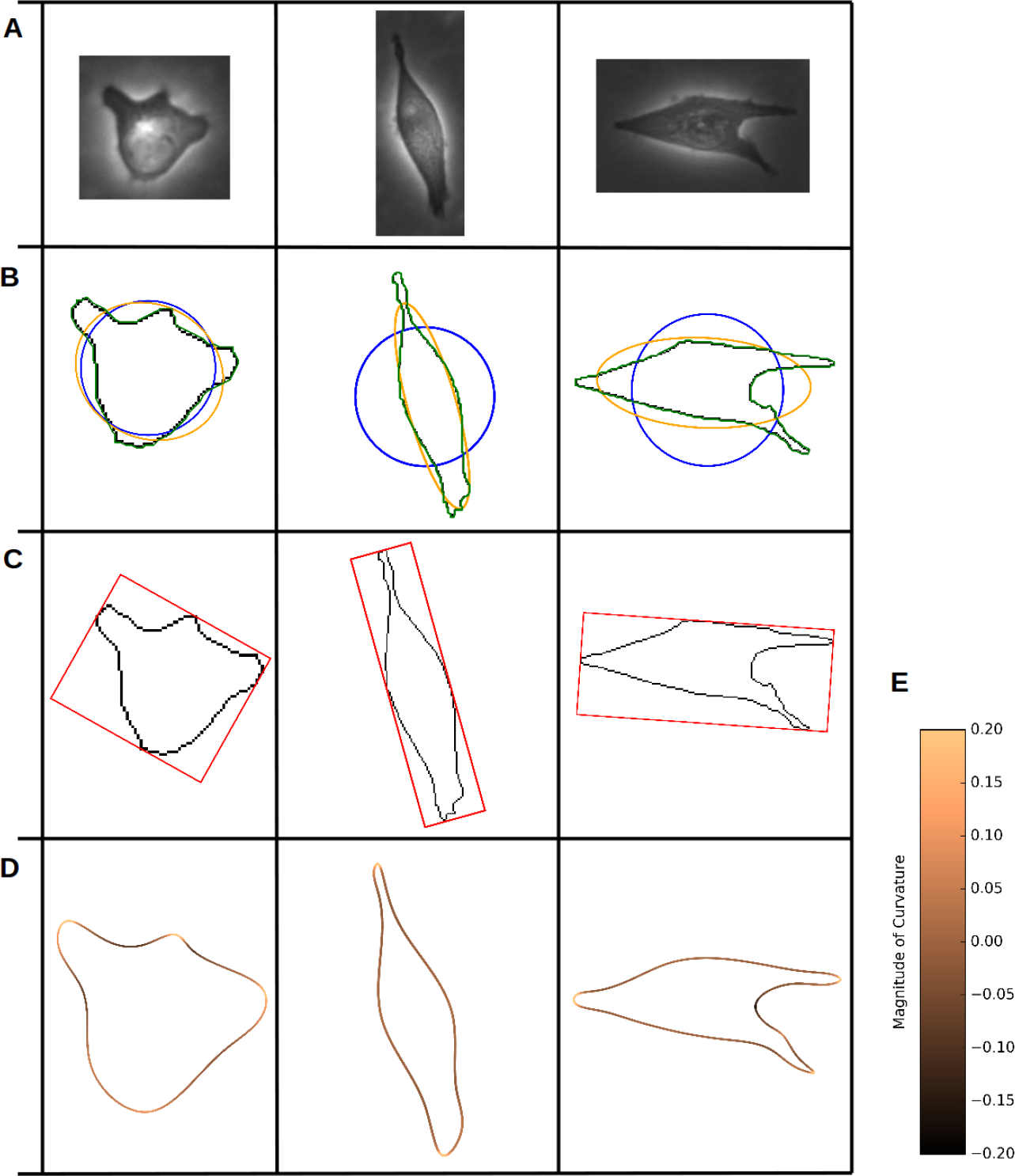
Geometrical fits and boundary interpolation. (A) Cropped images of cells from phase-contrast image, (B) Ellipse, circle and polygonal fit to segmented cell image, (C) Oriented rectangular fit, (D) Cubic spline fit along the boundary color coded by curvature, and (E) Color bar showing curvature values

**Figure 5.**
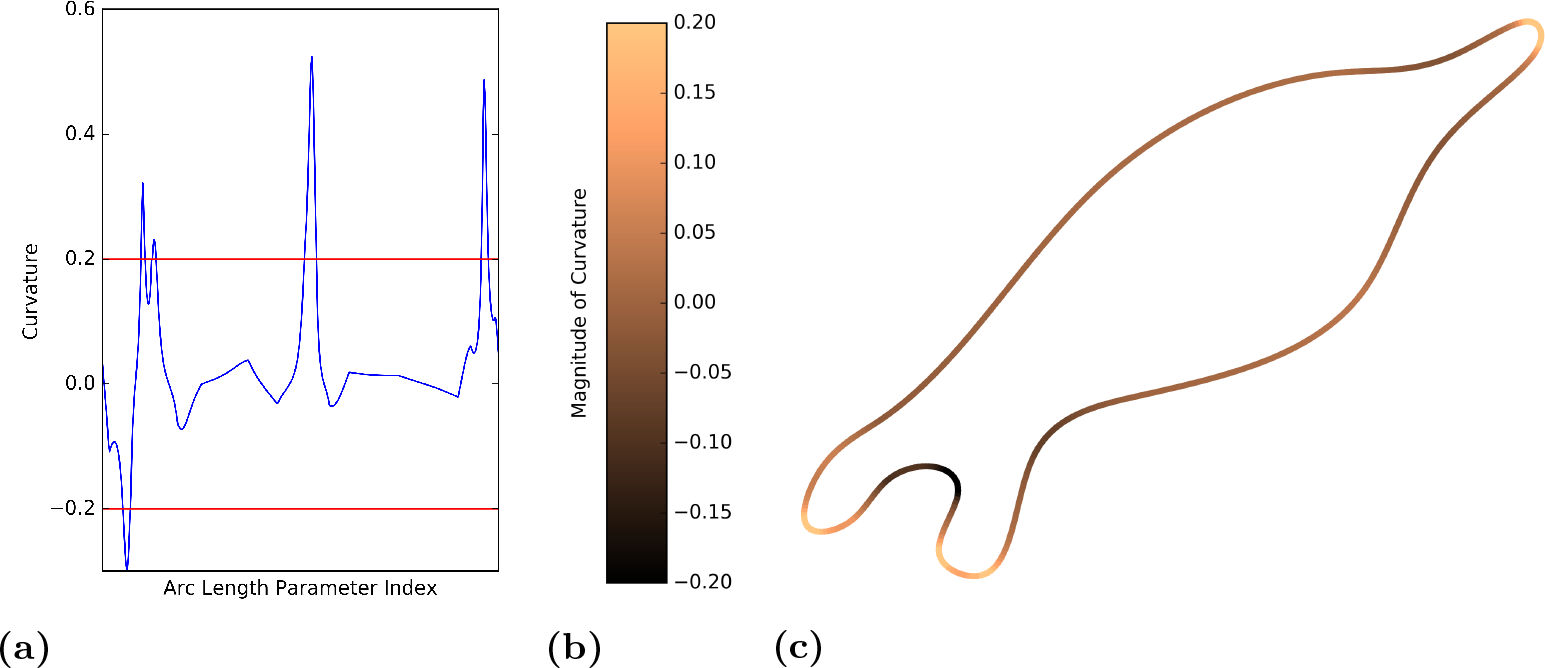
Computation of boundary features. (a) Curvature along the parametrized boundary obtained from cubic spline fit. (b) Colorbar for curvature values. (c) Cubic spline interpolation along the boundary color coded by curvature value. 3 protrusions and 1 indentation are detected in the cell shown above. Local maxima and minima in curvature with magnitude greater than 0.2 are classified as protrusions and indentations respectively. Multiple extrema within a neighborhood of 10 boundary pixels are treated as a single protrusion or indentation.

#### Ellipse and Circle Fitting

Using least-squares, an ellipse is fit to the cell outline so as to minimize the distance between closest points on the ellipse and the cell boundary. The optimal least squares solution can be computed directly without an iterative approach [31], as shown in S2 Appendix. The stable and robust fitting method returns the ellipse center, axes, and angle of rotation, *θ* = (*x*_*c*_, *y*_*c*_, *a*, *b*, *α*). In addition to parameters obtained from fitting, goodness of fit is estimated by calculating the variance [32]. Lengths of the two principal axes of the ellipse fit can be obtained from eigenvalues of the covariance matrix. Table 1 summarizes features obtained from ellipse and circle fitting.

**Table 1.** Features extracted from ellipse and circle fits of cell images

| Feature | Range | Description |
| --- | --- | --- |
| Ellipse Eccentricity | $[0, 1]$ | Close to 0, the ellipse is circular. Close to 1, it is elongated. |
| Ellipse Major Axis Length | $[0, \infty)$ | The length of the major axis of the ellipse fit. |
| Ellipse Minor Axis Length | $[0, \infty)$ | The length of the minor axis of the ellipse fit. |
| Ellipse Area | $[0, \infty)$ | Area of the ellipse fit. |
| Ellipse Perimeter | $[0, \infty)$ | Perimeter of the ellipse fit. |
| Ellipse Variance | $[0, 1]$ | A goodness of fit measure for the ellipse fit. |
| Circle Radius | $[0, \infty)$ | Radius of the circle fit. |
| Circle Area | $[0, \infty)$ | Area of the circle fit. |
| Circle Variance | $[0, 1]$ | A goodness of fit measure for the circle fit. |

#### Rectangle and Polygon Fitting

A rectangular fit for a cell consists of four functions with constraints, mandating an approach distinct from fitting conic sections. We use the procedure of Chaudhuri and Samal [33]: 1) finding the centroid of the object, 2) determining principal axes, 3) computing the upper and lower furthest edge points along the boundary, and finally, 4) finding the vertices of the bounding rectangle. Table 2 summarizes Feret diameters based on the rectangle fit. We use these to compute elongation, a non dimensional shape factor. Note the distinction between bounding box and the above rectangle fit. The edges of a bounding box are parallel to Cartesian axes whereas the major axes of the rectangular fit are aligned to the principal axes of the cell shape.

**Table 2.** Features extracted from bounding rectangle and polygonal fits

| Feature | Method | Description |
| --- | --- | --- |
| Maximum Feret Diameter | Rectangle fit | Longest distance between two parallel tangents on cell boundary |
| Minimum Feret Diameter | Rectangle fit | Shortest distance between two parallel tangents on cell boundary |
| Cell Area | Polygon fit, Pixel counting | Number of pixels inside segmented cell boundary. |
| Cell Perimeter | Polygon fit, 3-pixel Vector (3PV) Method | Estimate obtained from chain code for pixels on cell boundary. |

#### Polygon Fitting

A polygon fit along the cell boundary is computed using the 3-pixel vector (3PV) method described by Inoue and Kimura [24]. The 3PV method is designed for calculating the perimeter of low resolution raster objects, where counting the number of pixels at the boundary of the object results in inaccuracies. We modified the standard 3PV method to obtain vertices of the polygon fit while computing the cell perimeter. Details of our implementation are provided in S2 Appendix.

#### Cubic Spline Boundary Fitting and Curvature

Table 3 summarizes all boundary features extracted from the segmented cell shape. A cubic spline interpolation along the boundary of the segmented cell is computed using vertices of the polygon fit. Curvature values are calculated from the first and second derivative of the spline at 500 uniformly spaced points. Mean and standard deviation of curvature is included in the feature vector. To estimate the number of protrusions and indentations in the segmented cell boundary, the number of local maxima and local minima (with values above 0.2 and below 0.2 respectively) is also recorded. An additional constraint that no two maxima or minima can be located within a neighbourhood of 10 pixels is imposed to avoid small arbitrary fluctuations in curvature computation. Finally, the global extremum values are also included in the feature vector to distinguish cells that exhibit sharp filopodia. We observe that cells with circular morphology have positive curvature all along their boundary with zero protrusions or indentations. Elliptical cells tend to have two protrusions, situated at the extremities.

**Table 3.**
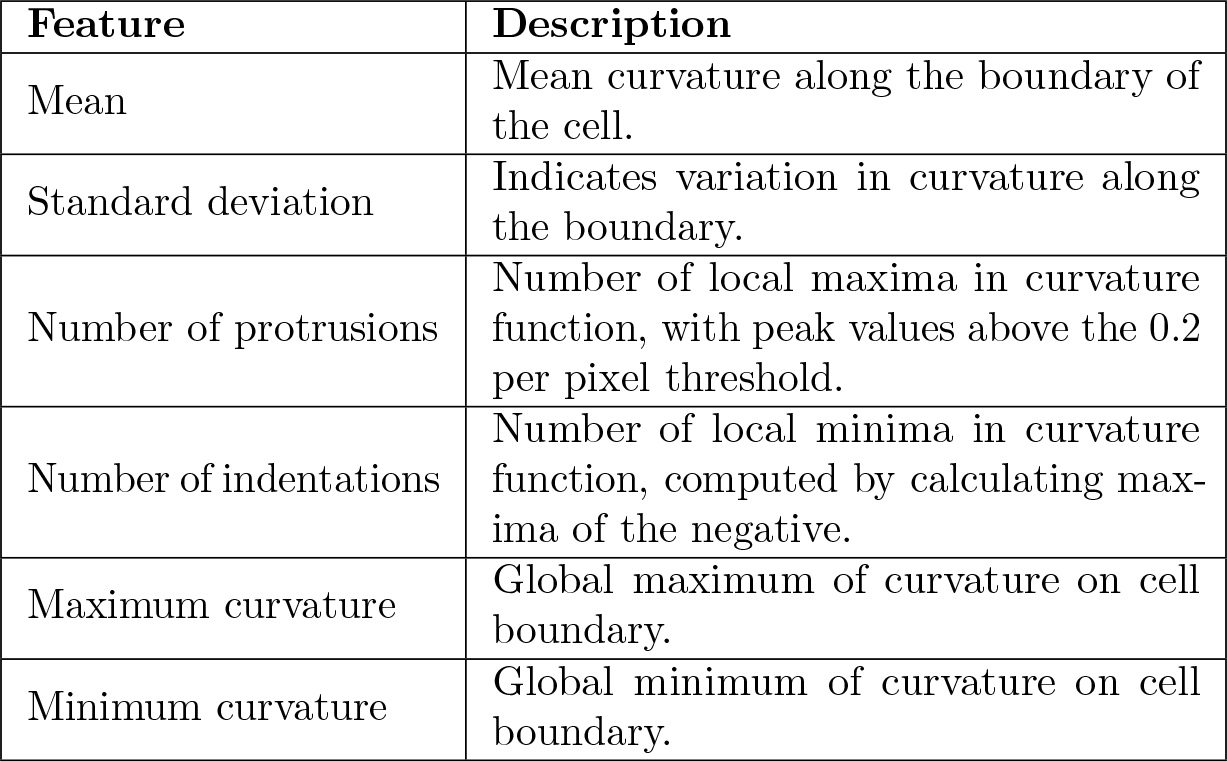
Features extracted from curvature of cubic spline fit

| Feature | Description |
| --- | --- |
| Mean | Mean curvature along the boundary of the cell. |
| Standard deviation | Indicates variation in curvature along the boundary. |
| Number of protrusions | Number of local maxima in curvature function, with peak values above the 0.2 per pixel threshold. |
| Number of indentations | Number of local minima in curvature function, computed by calculating maxima of the negative. |
| Maximum curvature | Global maximum of curvature on cell boundary. |
| Minimum curvature | Global minimum of curvature on cell boundary. |

#### Shape Factors

The previous section described features obtained by fitting various geometries to a cell boundary. Shape factors are non-dimensional quantities that are computed by counting pixels in a segmented cell image, and its convex hull, bounding box and oriented rectangular fit. Shape factors are widely used to classify particulate matter [34, 35] and often used as part of feature vectors designed to classify cell shapes [36, 37].

In Table 4, we list the non-dimensional shape factors included in our feature vector and formulas for their computation. These factors include *extent*, *solidity*, *compactness*, *elongation*, *circularity*, and *convexity*. More details about these geometric measures are provided in S2 Appendix.

**Table 4.** List of non-dimensional shape factors

| Feature | Range | Equation | Description |
| --- | --- | --- | --- |
| Extent | [0, 1] | $\frac{A_{\text{cell}}}{A_{\text{bounding box}}}$ | Ratio of pixels belonging to segmented cell to pixels in the bounding box. |
| Solidity | [0, 1] | $\frac{A_{\text{cell}}}{A_{\text{convex}}}$ | Ratio of pixels belonging to segmented cell to pixels in the convex hull. |
| Compactness | [0, 1] | $\frac{\sqrt{\frac{4(A_{\text{cell}})}{\pi}}}{\text{Max. Diameter}}$ | Ratio of circular equivalent diameter to maximum Feret diameter. |
| Elongation | [0, 1] | $1 - \frac{\text{Min. Diameter}}{\text{Max. Diameter}}$ | 1 - Aspect Ratio. Close to 1 for elongated cells and close to 0 for circular cells. |
| Circularity | [0, 1] | $\sqrt{\frac{4\pi A_{\text{cell}}}{P_{\text{cell}}^2}}$ | Degree of resemblance to a circle. |
| Convexity | [0, 1] | $\frac{P_{\text{convex hull}}}{P_{\text{cell}}}$ | Ratio of the convex hull perimeter to the cell perimeter. |

Briefly, the *extent* is the fraction of the bounding box area taken up by the cell, whereas *solidity* is the fraction of the convex hull occupied by the cell. For the MIA PaCa-2 pancreatic cancer data set, rounded cells typically have solidity values that approach unity. *Compactness* is the ratio of the diameter of a circle (with the same area as the cell) to the major axis of the rectangle fit whereas *elongation* is the ratio (1-*d*_1_/*d*_2_), where *d*_*i*_ are the width and length of the rectangular fit respectively. Compactness is close to zero if a cell is elongated, whereas elongation is close to zero if a cell is circular. *Circularity* measures the degree of similarity to a circle, whereas *convexity* is the ratio of convex hull perimeter to segmented cell perimeter.

### Dimensionality Reduction

Our unsupervised classification relies on clustering algorithms based on some distance metric. It tends to perform poorly for high dimensional feature vectors, since in high-dimensional space, distances between pairs of points tend to converge to similar values. To prevent this “curse of dimensionality”, our clustering is performed on a low-dimensional data set after dimensionality reduction.

#### Principal Component Analysis (PCA)

Principal Component Analysis, briefly summarized in S3 Appendix, is a common method for dimensionality reduction. PCA transforms high dimensional data into a low dimensional subspace, whose basis consists of linear combinations of the high dimensional basis vectors. The PCA basis vectors (PCA1, PCA2, etc), termed principal components, correspond to directions with the greatest variance in the original dataset. That is, (PCA 1) has maximum variation, (PCA 2) has the second-most variation, etc. A plot of variance in data explained by each principal component versus the number of components is generally used to identify how many components are needed. The tradeoff between number of components and total variance in data captured by principal components is resolved by finding the “elbow” in this plot. Typically 2-3 components account for majority of the explained variance and clusters in the transformed data can be easily visualized. However, as we demonstrate, this was not the case for the data used in this paper, motivating other dimensionality reduction methods.

#### t-Distributed Stochastic Neighborhood Embedding (t-SNE)

t-Stochastic neighborhood embedding (t-SNE) is a non-linear embedding from high dimensional space (e.g. *R*^*n*^) to low dimensional space (in our case, *R*^2^) designed to preserve local structure in the data [38]. The map is created by iterated steps of gradient descent that minimize Kullback-Leibler divergence, a cost function that represents difference of similarities of pairs of points in *R*^*n*^ and similarities of their images in the lower dimension representation. (In original SNE, Gaussian distributions of point-distance probabilities were used for the similarity comparison, whereas t-SNE employs a Gaussian in *R*^*n*^ and a Student-t distribution in the image space). An intuitive description of the iteration step was provided in [38]: image points in 2D move as if “connected by springs”, with “spring stiffness” corresponding to the deviation of similarities between the neighbor points in the data and in their images. t-SNE modified and generalized SNE to more strongly repel points that are “dissimilar” in the data, putting them far away in the 2D image, while keeping similar points close to one another.

Unlike PCA, where an explicit linear transformation is set up, t-SNE is a nonlinear map that aims to preserve probabilities of points being close or far from one another. Because t-SNE normalizes over local density, it produces clusters in 2D that have similar sizes. t-SNE does not faithfully represent geometry and in some cases even blurs the topology of the original data, so that distances in the 2D t-SNE plot should not be over-interpreted. An informative review of such eccentricities, with interactive illustrations, is given in the website https://distill.pub/2016/misread-tsne/.

The t-SNE algorithm requires multiple input parameters (“perplexity”, “early exaggeration”, learning rate and number of iterations). The perplexity is effective number of neighbors of a data point, and typically ranges between 5 and 50, providing a weighting between the importance of local and global properties. Wattenberg et al. [39] provide an excellent summary of the effect of perplexity and motivate the need for experimentation with a range of values of this parameter (see also the aforementioned URL). Linderman et al. [40] discuss how to optimize the selection of the exaggeration parameter and the step size.

The learning rate parameter plays an important role in preventing the algorithm from getting stuck in a local minimum. Analysis of properties of t-SNE and its usefulness in data visualization is provided in [40, 41]. While t-SNE is less straightforward than PCA, we found that it produced better results for our data and feature vectors.

### Unsupervised Classification

Once a low-dimensional data set is designated, the final step is to infer relationships between the data points (unsupervised classification). Clustering algorithms are used to automatically group the data points (descriptors or features in the context of machine learning) corresponding to the cell images. Clustering methods typically require parameter optimization to maximize classification accuracy [42]. We experimented with a variety of clustering methods and parameter settings, including k-means algorithm and DBSCAN (Density-based spatial clustering of applications with noise), as reviewed in S4 Appendix. DBSCAN defines clusters based on the density of packing - associating closely-packed nearest neighbors into a given category [43]. We used the hierarchical variant, HDBSCAN [44] in the results described further on.

The k-means algorithm requires one parameter (*k*), which is the number of expected clusters in the data. In many cases, including ours, the true value of *k* is not known *a priori*, so some way to estimate this value is desirable. Rousseeuw described a heuristic using silhouette coefficients to identify the number of clusters in a given data set [45]. Points are clustered using k-means for various values of parameter *k*. Assuming that the algorithm converges and gives stable results, the silhouette score is computed by calculating the average of silhouette coefficients for all data points. The number of clusters in the data set, i.e. the optimal value for *k*, is one that maximizes the silhouette score. We provide details and examples of this method in S4 Appendix.

The DBSCAN algorithm distinguishes between core points, boundary points and noise points using two parameters, *ϵ* and MinPts. Core points have at least MinPts points within *ϵ* radius neighbourhood. A border point has fewer than MinPts points within its *ϵ*-neighbourhood. Points that are neither core points nor border points are noise points. The DBSCAN algorithm iterates over all points and assigns them to clusters based on their reachability from core points. Each cluster contains at least one core point. The algorithm is sensitive to parameter values and does not perform well when the data contains clusters of various densities. The HDBSCAN algorithm eliminates the need to specify the *ϵ* parameter by repeatedly running DBSCAN at various spatial scales corresponding to different values of *ϵ*.

## Results

We applied the methods described above to 40 phase-contrast images of the MIA PaCa-2 cancer cell line. By visual inspection, we found that 310 cells were correctly segmented.

A typical segmentation result is shown in Figure 6. As expected, we found that boundaries of cells that are closely packed were not resolved by the watershed algorithm (see Figure 6b) since cells sharing foreground markers are treated as one object. To segment closely packed cells, many iterations of erosion were needed (see mathematical morphology). This, however, leads to poor results for spindle-shaped cells with long thin “tails”, as multiple erosions shorten, split or remove such tails. We observed that the number of erosion iterations (a parameter we varied) affected the range of cell sizes and population density at which correct segmentations were obtained. Even for isolated cells, fine details of cell boundaries such as tiny protrusions are lost due to erosion during segmentation.

Once segmented, cells are each assigned a unique identification number (UID), that associates the cell with each of its representations, from original image, to final cluster membership. Our dataset consisted of 310 correctly segmented cells, a relatively small dataset in the context of machine learning. Each of these cells was associated with a 30 component feature vector for the unsupervised classification process. The feature vector consisted of the 7 Hu’s invariant moments, 6 shape factors, 11 geometrical features and 6 boundary features. Because our cells are not pre-identified by human experts, we have no “ground truth” against which to compare results. Hence, we use visual inspection for the validation step.

**Figure 6.**
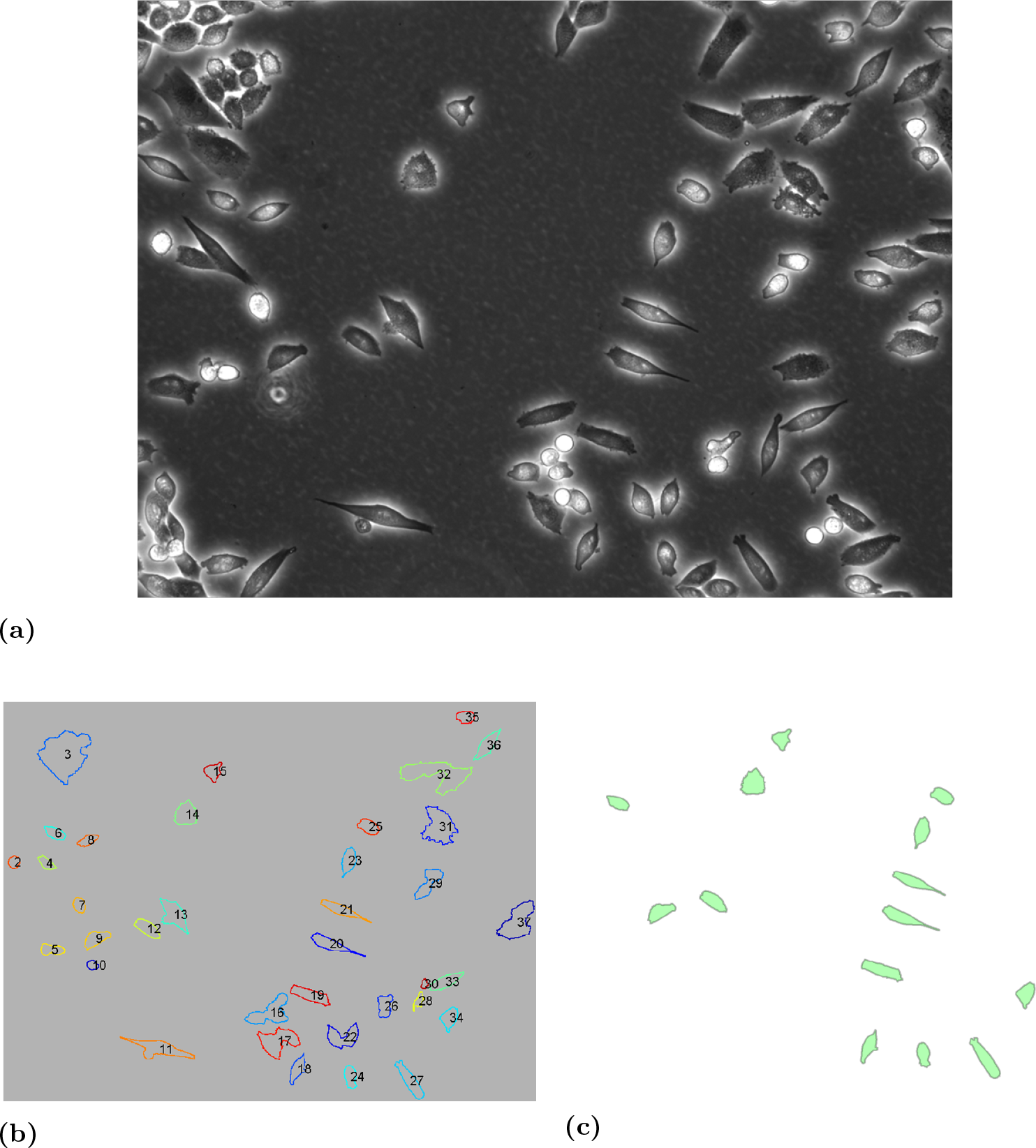
Cell segmentation. (a) MIA PaCa-2 image acquired using phase-contrast microscopy. (b) Boundaries (tagged with IDs) obtained using mathematical morphology and watershed segmentation. Each segmented boundary is assigned a unique ID. (c) Correctly segmented cells are serialized by manually selecting IDs (from Figure 6b). Features computed for IDs identified by the user are stored for further processing.

We first considered the K-means clustering algorithm on the entire feature vector of 30 features. To decide the optimal value of *K*, we computed silhouette scores for values of *K* ranging from 2 to 25. Since the cluster assignment obtained from K-means algorithm depends on the initialization of the the cluster centroids, the resulting silhouette scores also vary with each run of the algorithm. Averaging over 10 runs, we found that the silhouette score was maximized for *K* = 19, as shown in Fig. 7. This means that the K-means method specifies that the data consists of 19 clusters. The K-means algorithm is not capable of detecting outliers automatically, and determination of optimal value of *K* using silhouette score is often highly sensitive to the initial choice of centroids. We also considered using PCA for dimensionality reduction, as shown in Fig. 8. Note that clusters are not evident in 2 or 3 component PCA space, even though 3 components account for over 70% of the total explained variance in the dataset. This motivated us to switch the classification method.

Based on the suboptimal results obtained from PCA, we switched to a 2-component t-SNE followed by hierarchical clustering using HDBSCAN. All results described further on are based on this methodology. We classified cells using all 30 features, including the six shape factors, seven Hu’s moments, eleven geometrical features and six boundary features. We ran 1000 iterations of t-SNE with random initialization, perplexity value of 5, and learning rate of 200. Squared Euclidean distance was used to compute distance between features, which were standardized to have mean 0 and standard deviation of 1. The HDBSCAN algorithm was used to cluster feature points in the 2-component t-SNE space and identify outliers. The minimum points parameter was set equal to 4. We also tried multi-dimensional scaling (MDS), which produces a low dimensional representation of feature vectors with inter-point distances that are representative of distances in the higher dimensional feature space. However, similar to PCA, clusters were not evident in 2 and 3 dimensional MDS representation.

As shown in Figs. 9a and 9b, we found 27 clusters. There were also 19 outliers that did not get classified. Representative cells in each cluster are shown in Fig. 10. It is evident from Fig. 9a that some clusters (e.g. {1 and 2} as well as {14 - 21} are more closely related in the family tree of the HDBSCAN than others. As seen in Fig. 9b, these related clusters occupy nearby regions in the 2D plane of the t-SNE plot of Fig. 9b.

**Figure 7.**
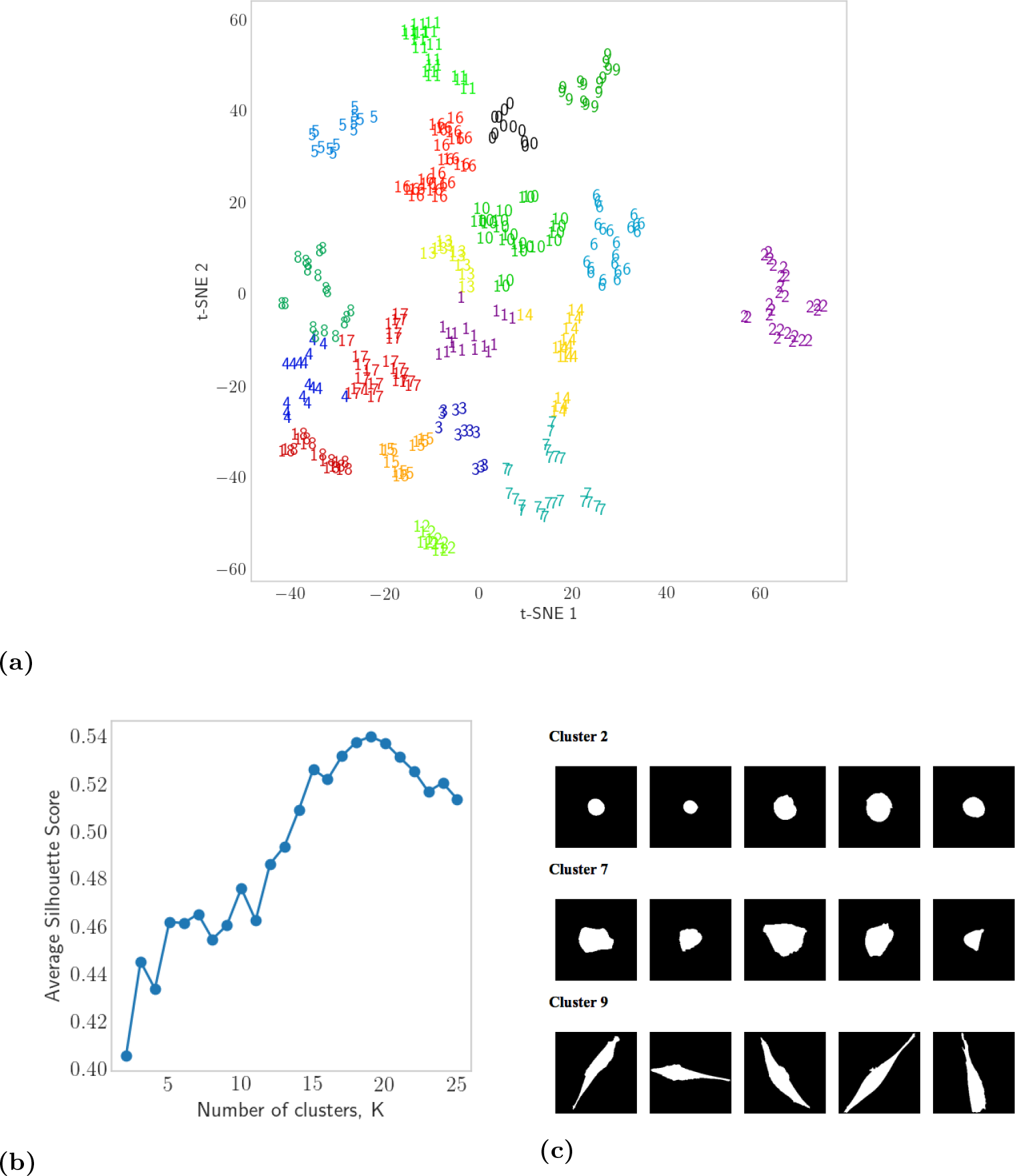
Classification using the K-means algorithm. (a) 19 clusters (labelled 0-18) obtained from 2-dimensional t-SNE features using the K-means algorithm. (b) The number of clusters, K. To find the optimal *K*, we perform clustering with a range of *K* values and find the value for which the silhouette score is maximized, here we find K = 19. Note that the outcome of K-means clustering depends on the initial choice of means. Consequently, we obtain different results with each run. (c) Selected clusters obtained using the K-means algorithm. Cells in each cluster have similar shapes.

**Figure 8.**
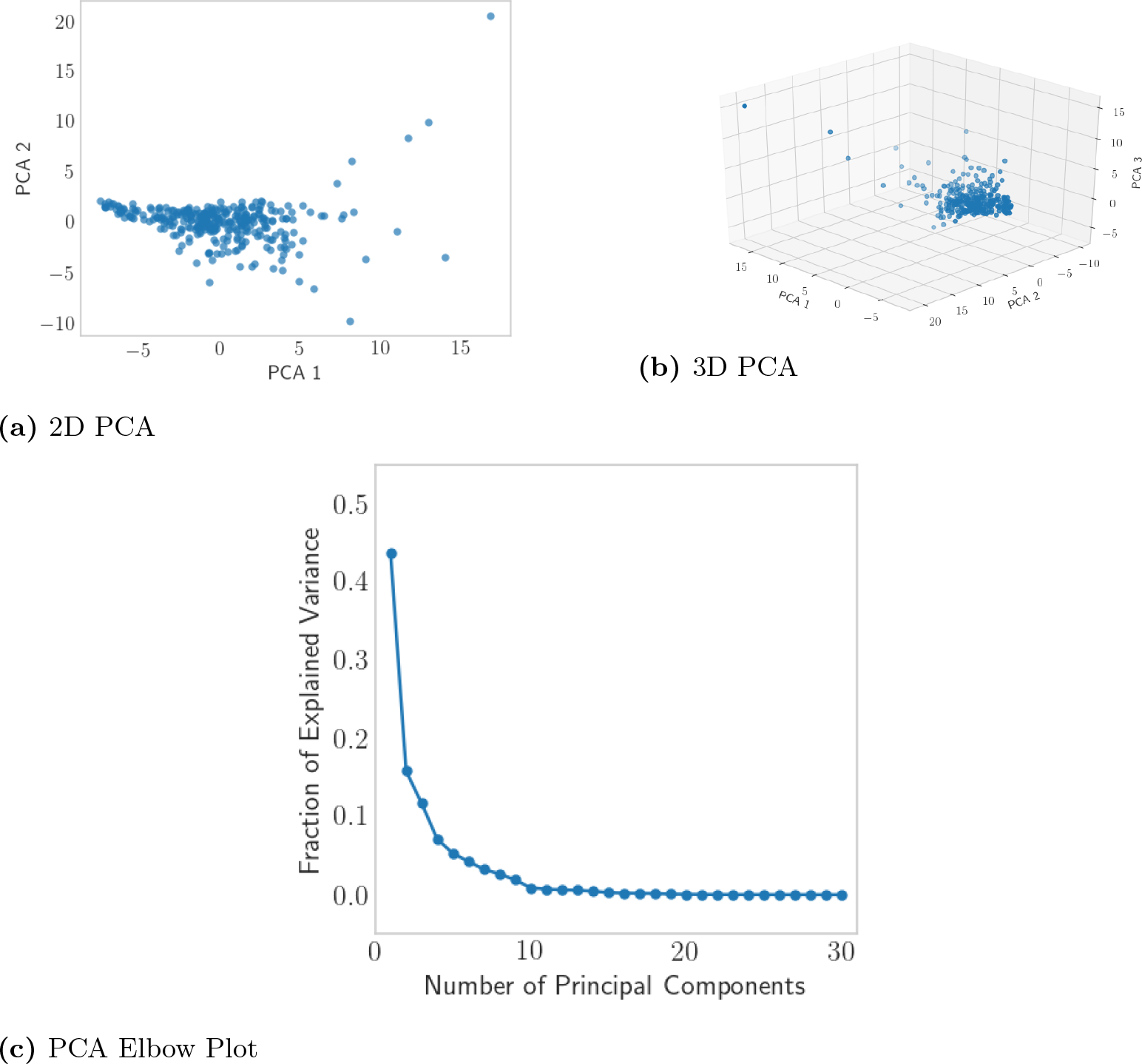
Dimensionality reduction using PCA. Although 3 principal components account for over 70% of the total variance in our data, we are not able to resolve clearly defined clusters when our features are embedded in 3-D PCA space. Instead, we see a single cluster and multiple outliers.

**Figure 9.**
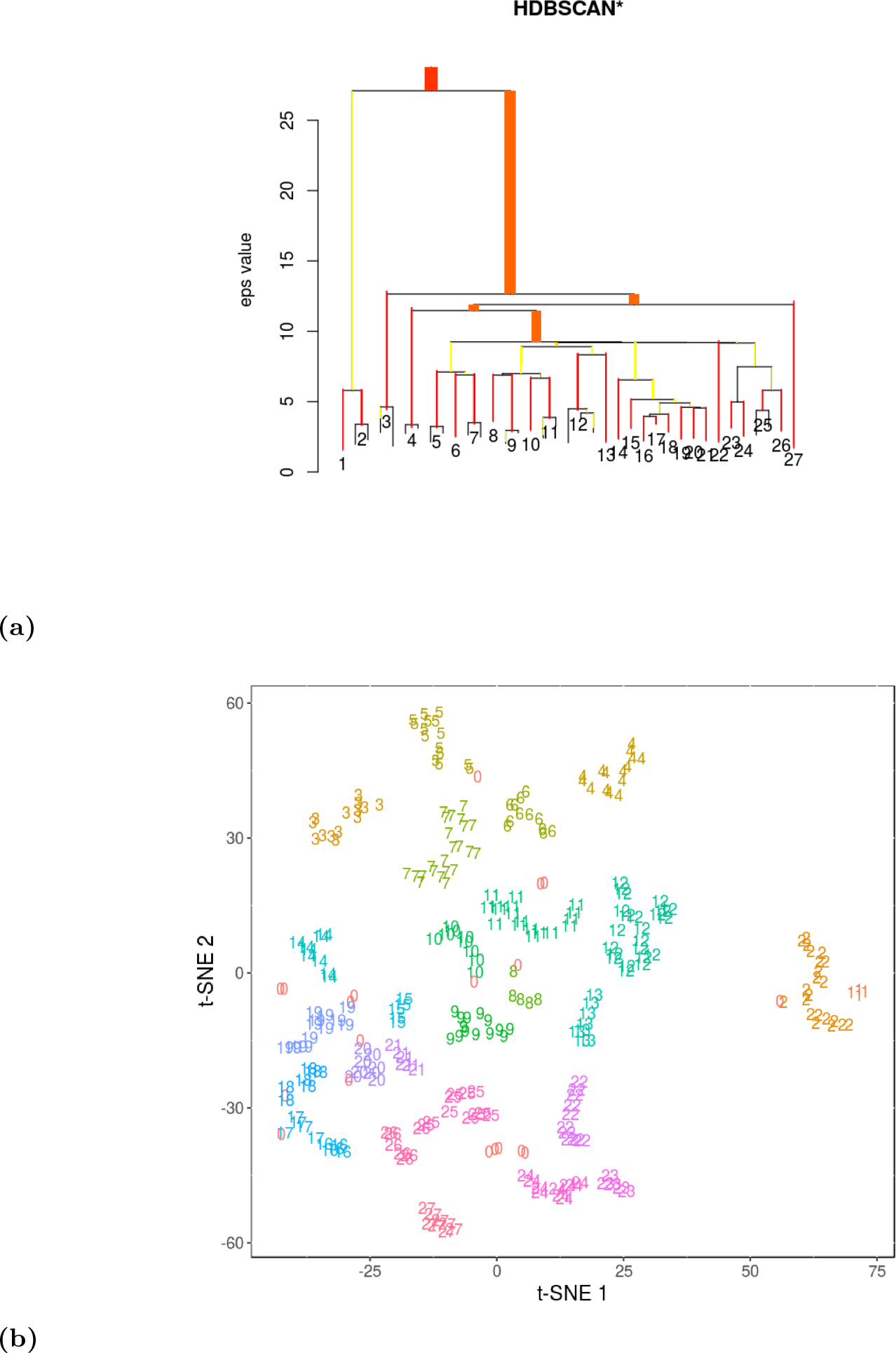
Clustering all features in a 2-component embedding. (a) Hierarchical clustering using HDBSCAN on all 30 cell features. The *ϵ* parameter in DBSCAN algorithm is varied to obtain density-based clustering hierarchy. HDBSCAN groups clusters into those that are more closely related along this “family tree” diagram. Cluster assignment in the 2D t-SNE plane. Cluster ID 0 indicates outliers that were not assigned to any of the 27 clusters identified. See Fig. 11 for a visualization of the cell shapes in a given cluster.

**Figure 10.**
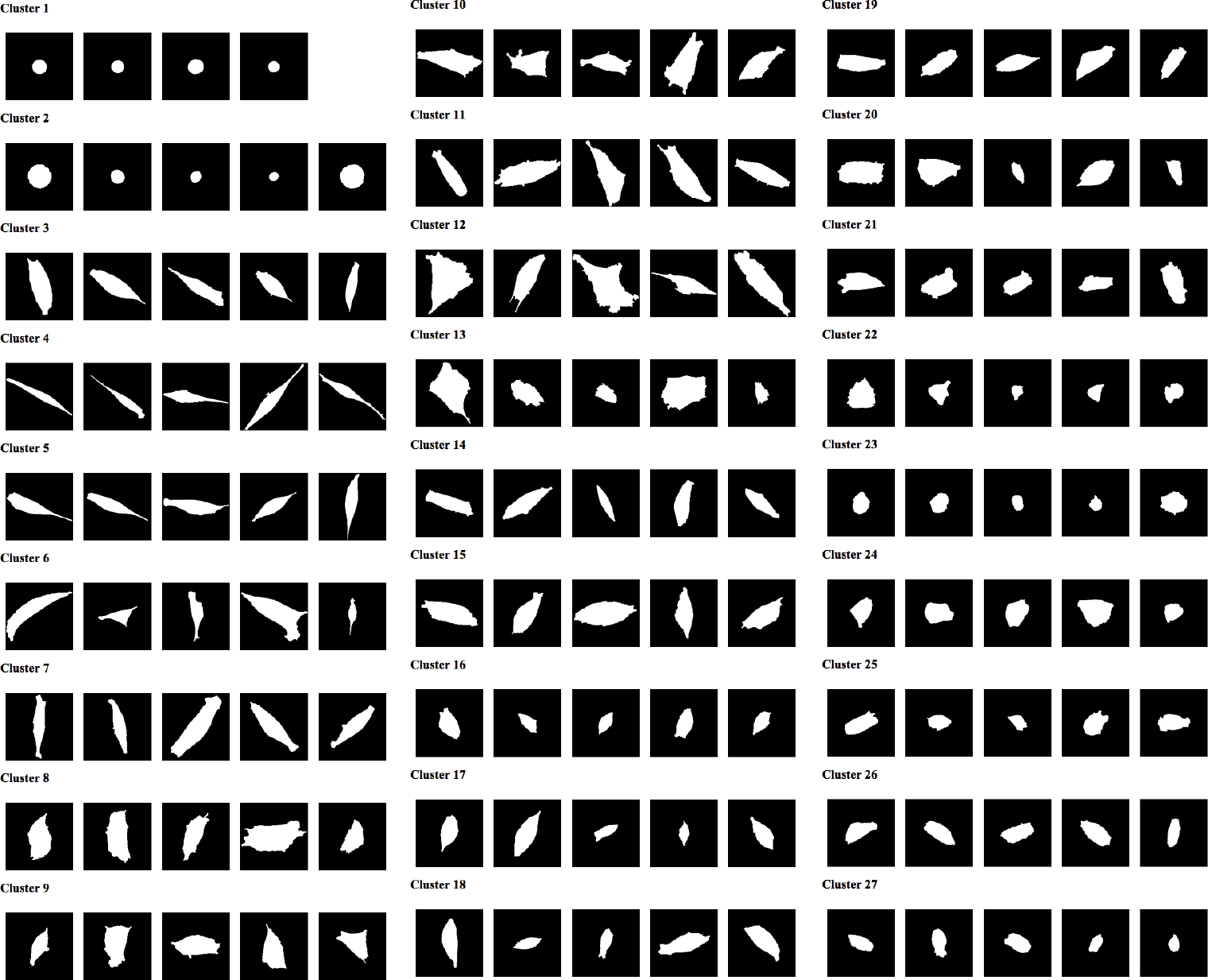
Clusters of cell shapes obtained by classifying shapes using all features.

In order to gain some insight into the clustering results, we manually created a composite diagram, Fig. 11 with HDBSCAN-related clusters grouped into ellipsoidal domains in the t-SNE 2-D plane. We also superimposed a representative cell shape on each cluster, to get an indication of how clustering was distinguishing between cells of different types, and how clusters were grouped in the t-SNE dimensionality reduction. We see from Fig. 11 that the 2D plane is roughly subdivided into circular cells (right), spindly and polarized cells (top), lens-shaped cells (left) and lumpy cells (bottom). The central region exhibits a larger number of irregularly spread out cells, some of which are fan-like, or polygonal. We also see that dimensionality reduction using t-SNE tends to preserve geometrical structure in data in the sense that similar shapes are grouped more closely in the 2-D t-SNE plane.

**Figure 11.**
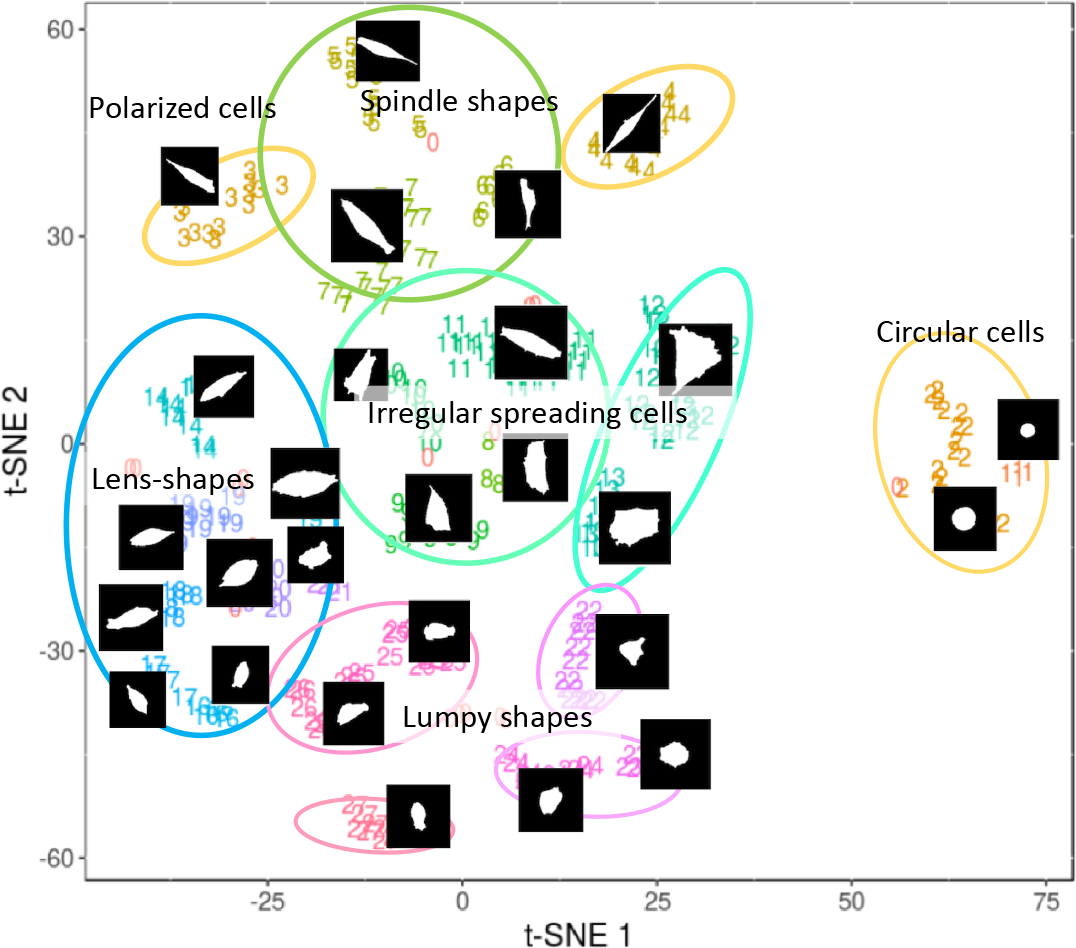
The 2D t-SNE plane, as in Fig. 9b, but with closely related clusters grouped, and with superimposed shapes of cells in each cluster. See also SM Fig. ?? for greater detail on the variety of cell shapes within each cluster.

### Comparisons with clustering using a subset of features

Next, we asked whether the entire 30-feature vector is essential for cell shape classification. To explore this question, we ran a limited number of tests with smaller subsets of the feature vector using HDBSCAN and 2-component t-SNE. These results are summarized in S5 Appendix and our conclusions are presented in Table 5. As shown in the table, the number of clusters detected ranged between 27 and 34, with between 19 and 14 outliers.

**Table 5.**
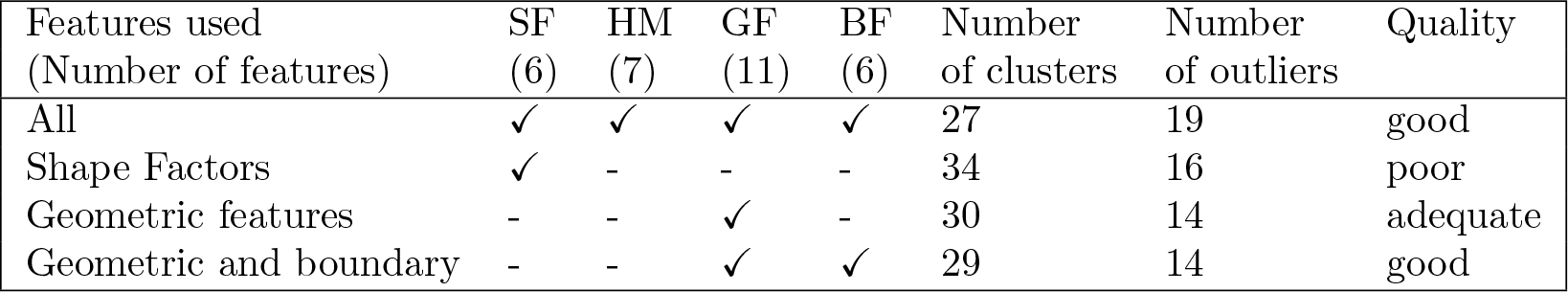
A comparison of the number of clusters and outliers found using subsets of the cell descriptors with 2-component t-SNE followed by hierarchical clustering using HDBSCAN. SF = shape factors, HM = Hu’s moments, GF = geometric features, BF = boundary features. See text for details on the quality of clusters.

| Features used<br>(Number of features) | SF<br>(6) | HM<br>(7) | GF<br>(11) | BF<br>(6) | Number<br>of clusters | Number<br>of outliers | Quality |
| --- | --- | --- | --- | --- | --- | --- | --- |
| All | ✓ | ✓ | ✓ | ✓ | 27 | 19 | good |
| Shape Factors | ✓ | - | - | - | 34 | 16 | poor |
| Geometric features | - | - | ✓ | - | 30 | 14 | adequate |
| Geometric and boundary | - | - | ✓ | ✓ | 29 | 14 | good |

In S5 Appendix we show several alternate classification results. Using the **six shape factors** alone, we found greater heterogeneity in cell size within clusters than with the classification based on the entire feature vector of 30 features. This is expected, since shape factors are invariant to scaling. If cells are classified based on **geometrical features**, which includes cell size, we observe that cells belonging to a cluster tend to be similar in size. Classification based on **geometrical and boundary features** are also demonstrated in S5 Appendix.

## Discussion

The main contribution of our paper is to develop a pipeline consisting of image analysis and unsupervised machine learning methods that is suitable for analyzing and classifying microscopy images that have no labels or annotations. The methodology we illustrate in this paper is particularly useful for finding patterns and relationships within large datasets where there is no knowledge about the basic classes of objects in advance. A second contribution is in extracting physically meaningful features from cell images, including cell shape and size, number of protrusions, and quantities that are less abstract than coefficients of orthogonal functions (Fourier or Zernike moments). The desirability of methods of classification based on physically meaningful descriptors has been pointed out recently in the literature [46].

In summary, we combined methods of cell segmentation as in [20, 22, 47] to first identify and segment cells. We then computed features similar to those used in [25, 26, 32, 34]. We added several of our own features (e.g. curvature of cell boundary) to compose a 30-dimensional feature vector. We experimented with PCA as in [17, 48] to reduce dimensionality, but found that t-SNE [38] results in a better embedding for cluster identification. While previous work employs a subset of these methods to perform cell classification, we believe that our pipeline is the first to combine these methods into a streamlined pipeline for unsupervised cell classification.

We have also shown that, with these features, we were able to achieve reasonable classification of cell shapes into categories that are clearly meaningful, consistent, and related. We showed that for the images and descriptors in our dataset, PCA does not work if we keep only the first two or three principal components, accounting for approximately 70% of the variance in the data. At that resolution, we can only separate outliers from the rest of the cells, as seen from Fig. 8. PCA would perform better if we keep a larger number of components (7 to 9, but then the results are not easily visualized). For this reason, dimensionality reduction using 2 component t-SNE, followed by clustering using HDBSCAN was the method of choice for us.

Comparing results obtained from a smaller subset of features (S5 Appendix) versus all features, we found less heterogeneity in cell size within each cluster if we use only the geometric features or geometric plus boundary features. This makes sense, since shape factors normalize the size of cells, preventing cell size from affecting the classification. Finally, we noted that for our data, geometric features by themselves produce reasonable qualitative results. In contrast, shape factors alone lead to poor results as these ignore differences in cell size. Similarly, boundary features alone (not shown in figures) are also inadequate, since these also fail to account for cell size. It is likely that data with larger variation in cell shape would require the combination of all 30 features to achieve good classification.

## Author Contributions

Conceptualization: HK, DB; Data curation: PD, DB; Formal analysis and Methodology: DB, DL, MZ, CT; Investigation and Project administration: DB; Funding acquisition and Resources: LEK, CR; Software: DB, DL, MZ; Writing: DB, LEK.

## Supporting information

**S1 Appendix. Mathematical morphology: erosion and dilation.** Definition of mathematical operations on binary images used to perform foreground detection.

**S2 Appendix. Feature computation.** Methodology and numerical methods used to compute Hu’s moments, elliptical and circular fits, polygonal fits and shape factors.

**S3 Appendix. Dimensionality reduction.** Description of Principal Component Analysis (PCA) and t-Distributed Stochastic Neighborhood Embedding (t-SNE).

**S4 Appendix. Unsupervised classification.** Clustering algorithms and silhouette score analysis.

**S5 Appendix. Classification using alternative feature vectors.** Results obtained using a low-dimensional feature vector containing only shape factors or geometrical features or Hu’s moments.

## Acknowledgments

We thank members of the Keshet-Feng and Roskelley research groups for discussions and helpful suggestions. The research reported here was supported by NSERC Discovery Grant (RGPIN-41870) awarded to Leah Edelstein-Keshet.

