## Supplementary material for "A methodology for morphological feature extraction and unsupervised cell classification": S1 Appendix

### Supplementary Appendix 1: A methodology for morphological feature extraction and unsupervised cell classification

#### Mathematical Morphology: Erosion and Dilation

Here we define mathematical morphology operations on binary images used to produce Fig. 2(c) in the main text.

Erosion of image $A$ by structural element (abbreviated "strel") $B$, resulting in image $E$ is defined as:

$$A \ominus B = \{z \in E | B_z \subseteq A\},$$

where $B_z$ is the translation of $B$ by the vector $z$:

$$B_z = \{b + z | b \in B\}, \forall z \in E.$$

In practice, erosion leads to shrinking or thinning of the binary image. Dilation of image $A$ by structural element $B$, resulting in image $E$ is defined as:

$$A \oplus B = \bigcup_{z \in B} A_z,$$

where $A_z$ is the translation of $A$ by the vector $z$:

$$A_z = \{a + z | a \in A\}.$$

Dilation is used to grow or thicken regions in a binary image. All other morphological operations can be defined by composing erosions and dilations.
