## Supplementary material for "A methodology for morphological feature extraction and unsupervised cell classification": S2 Appendix

### Supplementary Appendix 2: A methodology for morphological feature extraction and unsupervised cell classification

#### Feature Computation

##### Hu's Moment Invariants

The mathematical concept of moments can be used to quantify properties of an image as a whole. Since we are interested in shapes of cells in microscopy images, we use moments that preserve translational as well as rotational invariance.

Rotational invariance requires a nonlinear transformation that is not trivial to compute. Hu derived these nonlinear expressions from normalized central moments up to order three using algebraic invariants [1]. Hu's moment invariants are widely used for translation, scaling and rotation invariant pattern recognition, including recognition of typed English language characters [2]. Typically a feature vector for image classification is comprised of seven invariants:

$$\begin{aligned}
\phi_1 &= \eta_{20} + \eta_{02}, \\
\phi_2 &= (\eta_{20} - \eta_{02})^2 + 4\eta_{11}^2, \\
\phi_3 &= (\eta_{30} - 3\eta_{12})^2 + (3\eta_{21} - \eta_{03})^2, \\
\phi_4 &= (\eta_{30} + \eta_{12})^2 + (\eta_{21} + \eta_{03})^2,
\end{aligned}$$

$$\begin{aligned}
\phi_5 = {} & (\eta_{30} - 3\eta_{12})(\eta_{30} + \eta_{12})[(\eta_{30} + \eta_{12})^2 - 3(\eta_{21} + \eta_{03})^2] \\
& +(3\eta_{21} - \eta_{03})(\eta_{21} + \eta_{03})[3(\eta_{30} + \eta_{12})^2 - (\eta_{21} + \eta_{03})^2],
\end{aligned}$$

$$\phi_6 = (\eta_{20} - \eta_{02})[(\eta_{30} + \eta_{12})^2 - (\eta_{21} + \eta_{03})^2] + 4\eta_{11}(\eta_{30} + \eta_{12})(\eta_{21} + \eta_{03}),$$

$$\begin{aligned}
\phi_7 = {} & (3\eta_{21} - \eta_{03})(\eta_{30} + \eta_{12})[(\eta_{30} + \eta_{12})^2 - 3(\eta_{21} + \eta_{03})^2] \\
& -(\eta_{30} - 3\eta_{12})(\eta_{21} + \eta_{03})[3(\eta_{30} + \eta_{12})^2 - (\eta_{21} + \eta_{03})^2].
\end{aligned}$$

Note that $\phi_7$ has the additional property of being skew invariant and therefore can be used to distinguish between mirror images.

##### Feature Computation Using Geometrical Fits

In this section, we describe the numerical procedures for obtaining circle, ellipse and polygon fits for segmented cell shapes.

##### Ellipse and Circle Fitting

Consider an ellipse centered at $(x_c, y_c)$ and rotated by angle $\alpha$. Let $(x_t, y_t)$ be the closest point on the ellipse to boundary point $(x_i, y_i)$. Then the shortest distance $D_i$ from the boundary point to the ellipse is given by:

$$x_t = x_c + a\cos(\alpha)\cos(t) - b\sin(\alpha)\sin(t),$$

$$y_t = y_c + a\sin(\alpha)\cos(t) + b\cos(\alpha)\sin(t),$$
$$D_i = \sqrt{(x_i - x_t)^2 + (y_i - y_t)^2}.$$

In this case, the optimal least squares solution does not require iteration [3], and produces $\theta = (x_c, y_c, a, b, \alpha)$. In addition to parameters obtained from fitting, goodness of fit is estimated by calculating its variance as follows [4]:

Suppose $(\bar{x}, \bar{y})$ is the centroid and $(x_i, y_i)_{i=1}^{N}$ are the boundary points on the contour of the shape that is being fitted. Then covariance matrix of the contour is:

$$C = \frac{1}{N}\sum_{i=1}^{N} V_i V_i^T = \begin{pmatrix} c_{xx} & c_{xy} \\ c_{yx} & c_{yy} \end{pmatrix},$$

where

$$V_i = \begin{pmatrix} x_i - \bar{x} \\ y_i - \bar{y} \end{pmatrix},$$

and,

$$c_{xx} = \frac{1}{N}\sum_{i=1}^{N}(x_i - \bar{x})^2,$$
$$c_{xy} = \frac{1}{N}\sum_{i=1}^{N}(x_i - \bar{x})(y_i - \bar{y}),$$
$$c_{yx} = \frac{1}{N}\sum_{i=1}^{N}(y_i - \bar{y})(x_i - \bar{x}),$$
$$c_{yy} = \frac{1}{N}\sum_{i=1}^{N}(y_i - \bar{y})^2.$$

Lengths of the two principal axes of the ellipse fit can be obtained by calculating eigenvalues of the covariance matrix:

$$det(C - \lambda_{1,2} I) = 0,$$
$$\lambda_1 = \frac{1}{2}[c_{xx} + c_{yy} + \sqrt{(c_{xx} + c_{yy})^2 - 4(c_{xx}c_{yy} - c_{xy}^2)}],$$
$$\lambda_2 = \frac{1}{2}[c_{xx} + c_{yy} - \sqrt{(c_{xx} + c_{yy})^2 - 4(c_{xx}c_{yy} - c_{xy}^2)}],$$
$$\text{Ellipse eccentricity, } e = \frac{\lambda_2}{\lambda_1}.$$

Variance is the standard deviation of radial distance from the centroid to the boundary points divided by the mean. Variance close to zero indicates a good fit.

$$\text{Variance of fit} = \frac{\sigma_R}{\mu_R},$$

where,

$$\mu_R = \frac{1}{N}\sum_{i=1}^{N} d_i,$$

$$\sigma_R = \sqrt{\frac{1}{N}\sum_{i=1}^{N}(d_i - \mu_R)^2},$$

$$\text{and} \quad d_i = \sqrt{V_i^T C^{-1} V_i}.$$

##### Polygon Fitting

A polygon fit along the cell boundary is computed using the 3-pixel vector (3PV) method described by Inoue and Kimura [5]. The 3PV method is designed for calculating the perimeter of low resolution raster objects, where counting the number of pixels at the boundary of the object results in inaccuracies. Starting from an arbitrary location, adjacent boundary pixels are assembled in an ordered set. Each element in the set is an ensemble of three adjacent pixels enumerated in counterclockwise order. The spatial configuration of each 3-pixel ensemble is specified using a pair of integers from 0 to 7. The integers specify the direction of counterclockwise travel between consecutive pixels. Integers 0, 2, 4, 6, represent east, south, west and north directions respectively, as shown in Figure 1. Similarly, integers 1, 3, 5, 7, are used to encode southeast, southwest, northwest and northeast directions respectively for diagonal placement of pixels. Therefore, a 3-pixel ensemble corresponding to "L" shape (Figure 1) is represented by integers $[4, 6]$ indicating west and north direction of travel in counterclockwise manner. The ordered set of integer representations for 3-pixel ensembles starting from the arbitrary location is defined as the chain code of the object boundary.

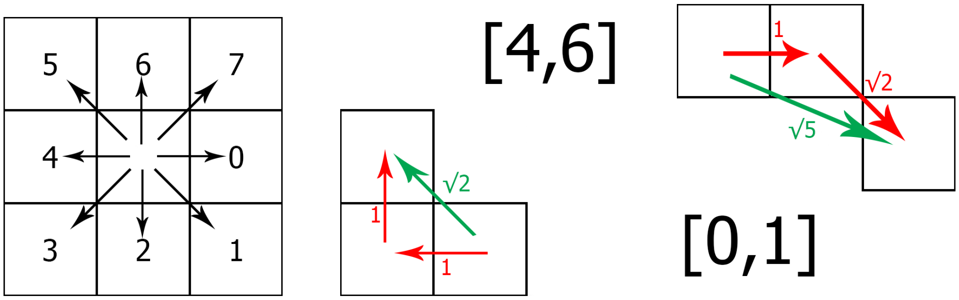


**Figure 1.** Computing chain code from boundary pixels
(a) Reference diagram from computing chain code. (b) An example of 3-pixel configuration corresponding to $\sqrt{2}$ length. (c) An example of 3-pixel configuration corresponding to $\sqrt{5}$ length.

As part of geometrical feature extraction, 3PV method is used to compute the perimeter of segmented cell images from the chain code representation of the cell boundary. An adjacent pair of elements in the chain code is referred to as a chain pair. Inoue and Kimura [5] specify corrections to typical perimeter calculation (the so-called $1, \sqrt{2}, \sqrt{5}$ method, illustrated in Figure 1, for computing distances for straight, diagonal, and straight followed by diagonal placement of pixels respectively) for all possible combinations of chain pairs. A set of vectors consisting of pairs of points along the cell boundary (called 3-pixel vectors, since they are derived from chain pairs corresponding to 3-pixel ensembles) is computed where the sum of lengths of these vectors provides an accurate estimate of the perimeter of the cell boundary. With minor adjustments (to account for cases where adjacent vectors do not align head to tail) as shown in Figure 2, the 3-pixel vector is used to obtain a polygonal fit to the cell geometry.

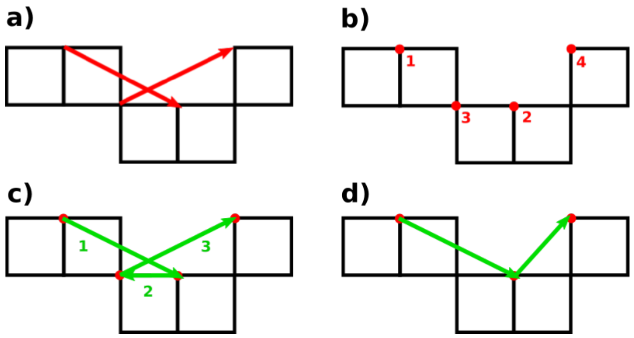


**Figure 2.** Computation of polygonal fit from chain code
(a) 3-pixel vectors obtained from chain code $[0, 1, 7]$ corresponding to east, southeast and northeast direction of travel between two adjacent 3-pixel ensembles on the cell boundary. (b) Ordered set of points along the cell boundary corresponding to the 3-pixel vectors. (c) Connecting points in order results in incorrect polygon segment as 3-pixel vectors are not aligned head to tail. (d) A vertex is removed to correct the boundary segment.

#### Shape Factors

We provide descriptions of the shape factors in this section.

##### Extent

The ratio of the number of pixels belonging to a segmented cell to the number of pixels in its bounding box is defined as the extent. The bounding box spans horizontally from the leftmost pixel to the rightmost pixel and vertically from the topmost pixel to the bottommost pixel. Extent is close to zero if a cell is elongated and close to unity if the cell is uniformly spread out.

##### Solidity

Solidity is a measurement of the overall concavity of an object. It is defined as the ratio of the number of pixels belonging to the segmented cell to the number of pixels in its convex hull. As cell shape deforms from a convex polygon or circle to a more elliptical or protrusive shape, its convex hull area increases compared to the cell area and solidity correspondingly decreases. For the MIA PaCa-2 pancreatic cancer data set, rounded cells typically have solidity values that approach unity.

##### Compactness

Compactness is defined as the ratio of the circular equivalent diameter to the maximum Feret diameter obtained from the bounding rectangular fit:

$$\text{Compactness} = \frac{\sqrt{\frac{4(A_{\text{cell}})}{\pi}}}{\text{Max. Feret diameter}}$$

The circular equivalent diameter, also known as area-equivalent diameter, is defined as the diameter of a circle with the same area as the object. Like extent, compactness is close to zero if a cell is elongated or ‘I’ shaped.

##### Elongation

Elongation is defined as (1 − Aspect Ratio). Aspect ratio is obtained from the rectangle fit as the ratio of minimum to maximum Feret diameters. For elongated cells, maximum Feret diameter is much larger than minimum Feret diameter, therefore their elongation is close to unity. Conversely, for circular cells, both diameters are roughly the same. Therefore the elongation of such cells is close to zero.

**Circularity**
Circularity measures the degree to which an object is similar to a circle:

$$\text{Circularity} = \sqrt{\frac{4\pi A_{\text{cell}}}{P_{\text{cell}}^2}}$$

It can be easily verified that circularity for a perfect circle is unity. Regular polygons approach a circle as their number of edges increases. It should be noted that a low value of circularity does not necessarily mean that the cell shape lacks rotational symmetry. Circularity close to zero typically indicates elongated or protrusive (e.g. starfish-like) morphology.

**Convexity**
Convexity is defined as the ratio of the convex hull perimeter and the actual perimeter of the object. It is close to zero for highly non-convex cell geometries and close to unity for epithelial-like cells with polygonal morphology (absent from MIA PaCa-2 data set) or circular cells.
