## Supplementary material for "A methodology for morphological feature extraction and unsupervised cell classification": S3 Appendix

### Supplementary Appendix 3: A methodology for morphological feature extraction and unsupervised cell classification

#### Dimensionality Reduction

Below, we compare two commonly used methods to obtain a low dimensional representation of high dimensional data.

##### Principal Component Analysis (PCA)

Principal Component Analysis is a mathematical technique that exploits variance in a given data set in order to make patterns in the data salient. It is commonly used in machine learning to visualize and manipulate high dimensional feature vectors. PCA transforms high dimensional data into a low dimensional subspace, where the basis for the low dimensional subspace is a linear combination of high dimensional basis vectors. The linearity assumption reduces the problem of finding an appropriate transformation to the problem of finding an appropriate projection. The linear combination of the original basis is determined in a manner such that the low dimensional basis vectors (also known as principal components) correspond to direction with the greatest variance in the data. In other words, PCA projects high dimensional feature vectors to a new (low dimensional) coordinate system, ensuring that the first axis in the new coordinate system (PCA 1) has maximum variation, the second axis (PCA 2) has the second-most variation, and so on.

Mathematically, the principal components are the eigenvectors of the covariance matrix of the original data set. These eigenvectors are orthogonal since the covariance matrix is symmetric. Reducing dimensionality of the feature space by projecting all feature vectors to a low dimensional space reduces the complexity of cluster identification and k-means clustering. The number of principal components (i.e. dimensionality of the transformed space) is chosen by plotting the variance in data explained by each principal component versus the number of components. Typically, the number of principal components is determined by finding an "elbow" in this plot. The elbow signifies the turning point where the trade-off between including additional variance is offset by the complexity of dealing with more components. Generally, most of the variance in the original data is explained by a small number of principal components. Sometimes 2 or 3 components are chosen for ease of plotting.

##### t-Distributed Stochastic Neighborhood Embedding (t-SNE)

t-SNE tends to preserve local structure in data, and so it is often used in place of PCA for dimensionality reduction. Unlike PCA, t-SNE is a non-parametric learning algorithm that handles non-linearity in the data very well. The embedding is learned in the process of moving data to the low dimensional space. Consequently, t-SNE does not provide a function for transforming data from the high dimensional space to the low dimensional space. Furthermore, the t-SNE algorithm requires multiple input parameters including perplexity, early exaggeration, learning rate and number of iterations. While default values of these parameters work well for widely publicized open data sets, the algorithm is sensitive to perplexity and learning rate parameters for features included in the MIA PaCa-2 data set. The perplexity parameter is similar to k in the k-nearest neighbors (KNN) classifier algorithm. It is used to build a nearest neighbor graph in the high dimensional feature space. The t-SNE model building process involves performing random walks on this feature graph. The learning rate parameter plays an important role in preventing the algorithm from getting stuck in a local minimum while minimizing the Kullback-Leibler divergence, a non-convex cost function. For more details on the t-SNE algorithm, please refer to Maaten and Hinton (2008) [1].

Shortcomings of t-SNE, include its stochastic nature (requiring multiple runs to ensure convergence), absence of parameter estimation techniques and lack of simplicity (compared to PCA where the linear transformation can be easily analyzed).
