## Supplementary material for "A methodology for morphological feature extraction and unsupervised cell classification": S4 Appendix

### Supplementary Appendix 4: A methodology for morphological feature extraction and unsupervised cell classification

#### Unsupervised Classification

Unsupervised classification refers to grouping of quantifiable objects by inferring relationships between these objects. Clustering algorithms are used to automatically group the data points (descriptors or features in the context of machine learning) corresponding to the objects of interest.

Irrespective of the method used for clustering the data, it is important to find parameters to optimize algorithm performance and ensure that the results obtained from the algorithm are meaningful. Many clustering algorithms are known to be very sensitive to their input parameters [1]. Measures for assessing the efficacy of clustering like Davies-Bouldin index and silhouette score (defined below) are useful for evaluating the results of clustering algorithms.

The k-means algorithm and DBSCAN are two widely used methods for clustering data in low dimensions. While the k-means algorithm requires one parameter ($k$), the DBSCAN algorithm requires two parameters (minPts, the minimum number of points in the neighborhood of a core point and neighborhood distance $\epsilon$) for distinguishing between core points, boundary points and non-reachable points.

A number of methods exist to estimate the value of parameter $k$ for k-Means clustering. We found four such methods that are widely used in literature: elbow heuristic, Bayesian information criterion (abbreviated BIC), silhouette analysis [2] and gap statistic [3]. In practice, using features computed from the MIA PaCa-2 data set, silhouette analysis (described below) performed best in terms of robustness and convergence.

#### Silhouette Score Analysis

Silhouette analysis is the study of the degree of separation between clusters of data points using silhouette coefficients. The silhouette coefficient for a given data point, $P$, is a measure that quantifies the degree to which the data point belongs to its assigned cluster, $C$. It is computed as follows. Let $a$ be the mean distance between point $P$ and every other point in its own cluster $C$. Let $b$ be the mean distance between $P$ and every point in the nearest neighboring cluster. Then, the silhouette coefficient for point $P$ is $(b - a)/\max(a, b)$. The silhouette coefficient ranges from -1 to 1. A coefficient value near 1 indicates that $P$ has undoubtedly been classified correctly, a value around 0 indicates that the clustering of $P$ has some ambiguity, and a value near -1 indicates it is likely that $P$ was classified incorrectly.

Rousseeuw described a heuristic using silhouette coefficients to identify the number of clusters in a given data set [2]. Points are clustered using k-means for various values of parameter $k$. Assuming that the algorithm converges and gives stable results, the silhouette score is computed by calculating the average of silhouette coefficients for all data points. The number of clusters in the data set, i.e. the optimal value for $k$, is one that maximizes the silhouette score. This technique is demonstrated using synthetically generated data in Figure 1. 10,000 data points corresponding to 10 clusters (1000 points per cluster) are generated by transforming and combining uniform random distributions (see Figure 1a). Figure 1b shows a plot of silhouette score computed for various values of parameter $k$. The most probable value of $k$ is automatically determined by finding the maximum in this plot. For the synthetic data set, the silhouette score is maximized at $k = 10$. Figure 1c shows the k-means clustering result (corresponding to $k = 10$), with points colored according to their cluster label. Sorted values of silhouette coefficients for individual data points (grouped by cluster label) are shown in Figure 1d, with the vertical red line depicting the overall silhouette score obtained by averaging.

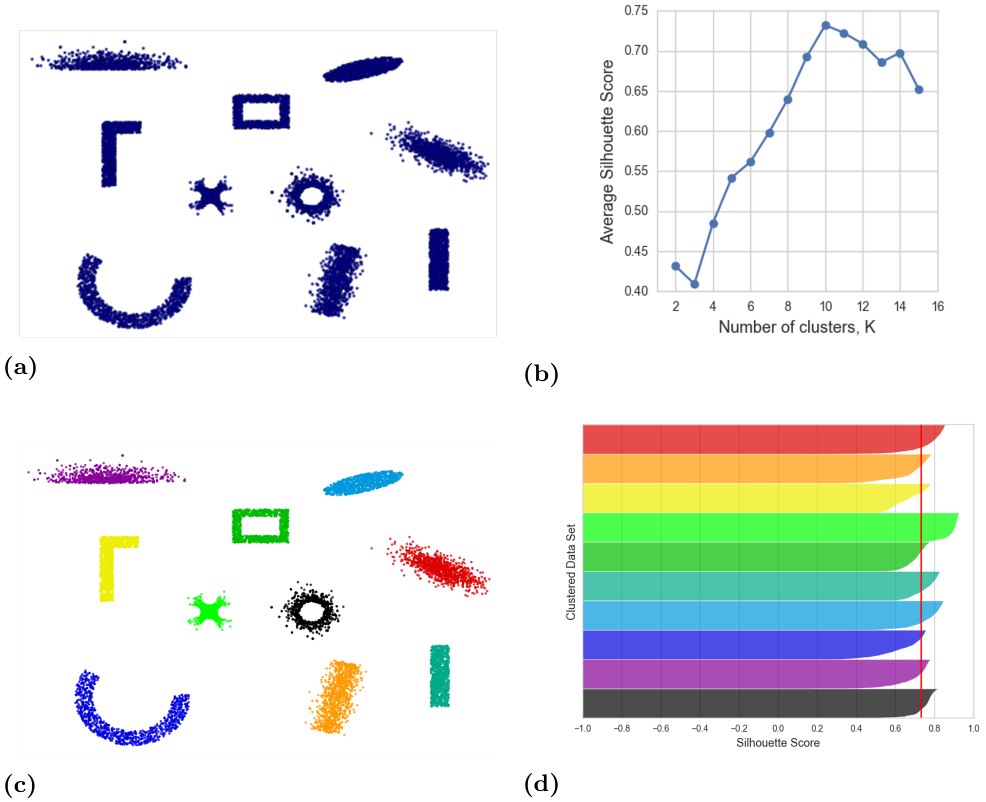


**Figure 1.** Clustering synthetic data using silhouette score analysis
(a) 10,000 synthetic data points corresponding to 10 user-defined cluster shapes. (b) Determining optimal $k$ automatically by computing average silhouette score. (c) Labeled data points (obtained by k-means) corresponding to 10 clusters. (d) Sorted silhouette scores for data points in each cluster.

#### References

1. Kovács F, Legány C, Babos A. Cluster validity measurement techniques. In: 6th International symposium of hungarian researchers on computational intelligence.

Citeseer; 2005.Available from: `http://citeseerx.ist.psu.edu/viewdoc/download?doi=10.1.1.100.2848&rep=rep1&type=pdf`.

2. Rousseeuw PJ. Silhouettes: a graphical aid to the interpretation and validation of cluster analysis. Journal of computational and applied mathematics. 1987;20:53–65.

3. Tibshirani R, Walther G, Hastie T. Estimating the number of clusters in a data set via the gap statistic. Journal of the Royal Statistical Society: Series B (Statistical Methodology). 2001;63(2):411–423. doi:10.1111/1467-9868.00293.
