## Supplementary material for "A methodology for morphological feature extraction and unsupervised cell classification": S5 Appendix

### Supplementary Appendix 5: A methodology for morphological feature extraction and unsupervised cell classification

#### Classification Using Alternative Feature Vectors

##### Alternate feature vectors

In this section we describe results that use the same methods, using HDBSCAN and 2-component t-SNE, but a smaller set of features.

**Classification using shape factors**: Here we used 6 shape factors: extent, solidity, compactness, elongation, convexity, circularity. We found 34 clusters. Results are shown in Fig 5. The collection of shapes of cells in the clusters is given in Fig. 1.

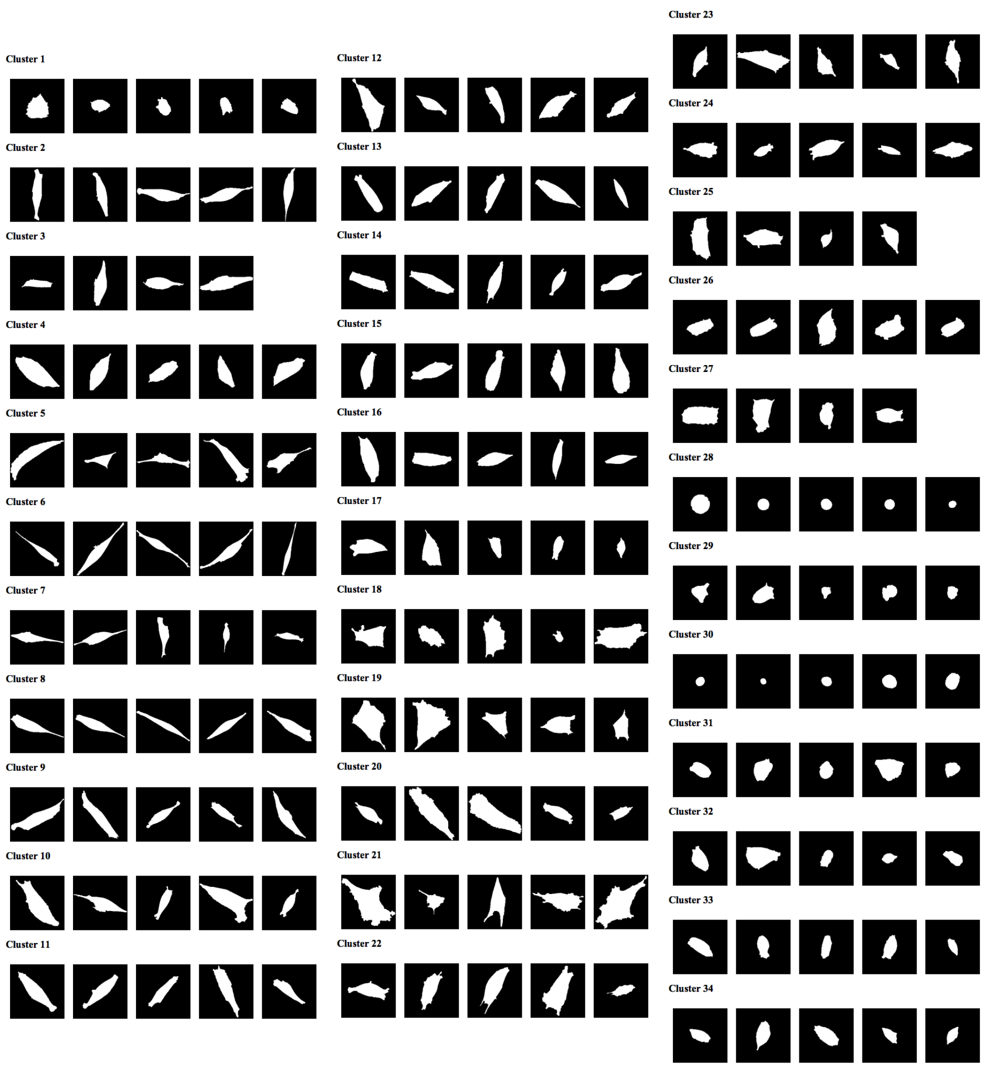


**Figure 1.** Clusters obtained by classifying cell shapes using only shape factors.

**Classification using geometrical features**: Here we used 11 features: polygon fit (2), ellipse fit (4), circle fit (3) and rectangular fit (2) We found 30 clusters. Results are shown in Fig 6. For purposes of comparison, we also show a composite image (Fig. 7) in which clusters that are closely related in the HDBSCAN "family tree" are grouped together and typical cell images superimposed. (Compare with a similar image in the main text where all features were included in the clustering.) We see that clustering using geometric features alone does a reasonably good job of separating out cell shapes, and that the 2D t-SNE plot groups similar clusters in nearby areas of the plane. Cells that are nearly circular are now at the bottom, with more polar and spindle shapes at the left, lens-shaped cells at the top, and lumpy cells on the left. The collection of shapes of cells in the clusters is given in Fig. 2.

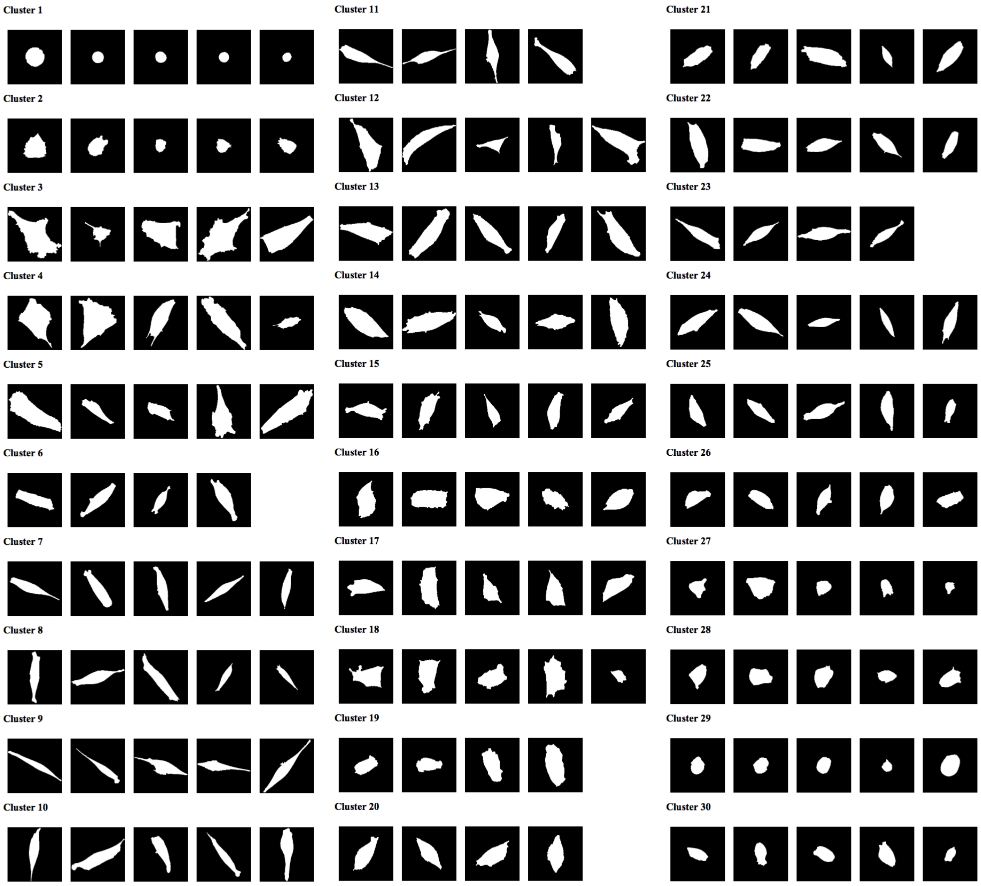


**Figure 2.** Clusters of cell shapes obtained by classifying shapes using only geometrical features.

**Classification using geometrical and boundary features**: We used 17 features: 11 geometrical features and 6 boundary features, including mean and standard deviation of curvature, number of protrusions and indentations, global maxima and minima of curvature. There were 29 clusters found. Results are shown in Fig 8. The shapes of cells in the clusters in shown in Fig. 3.

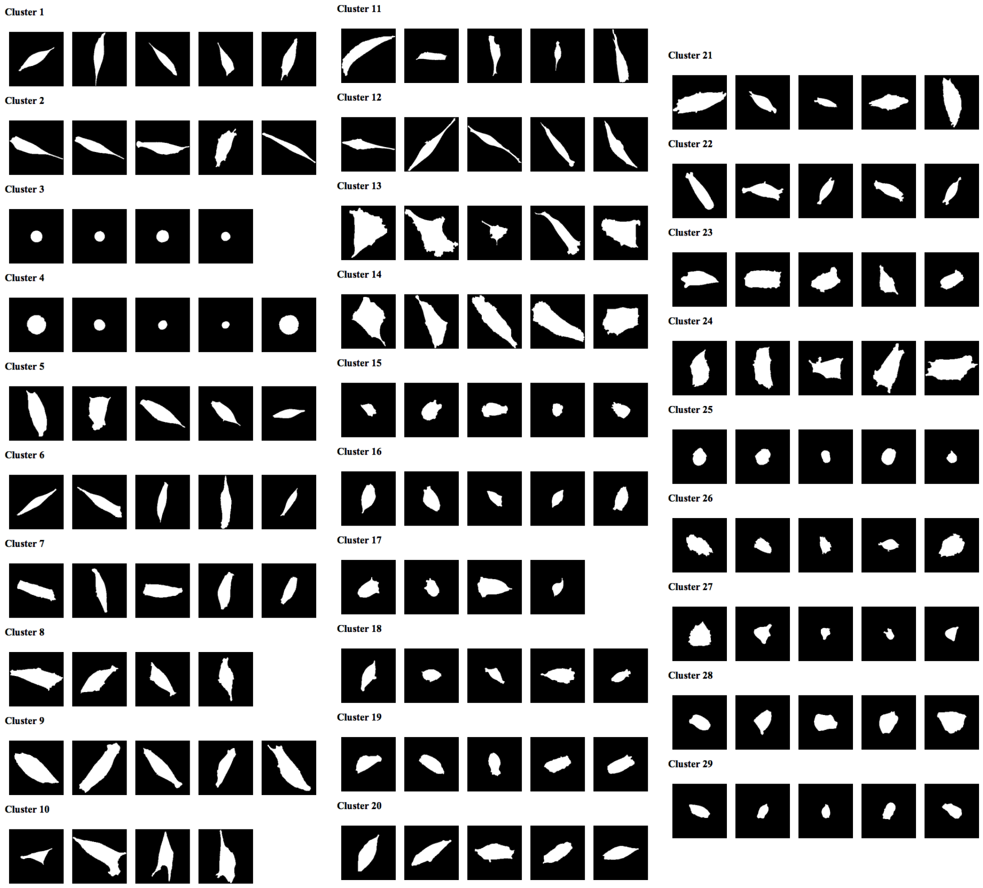


**Figure 3.** Clusters obtained by classifying cell shapes using geometrical and boundary features.

**Classification using Hu's moments**: We used 7 Hu's moments described in the methodology section. 33 clusters were found using t-SNE and HDBSCAN. Results are shown in Fig 9. The shapes of cells in the clusters in shown in Fig. 4.

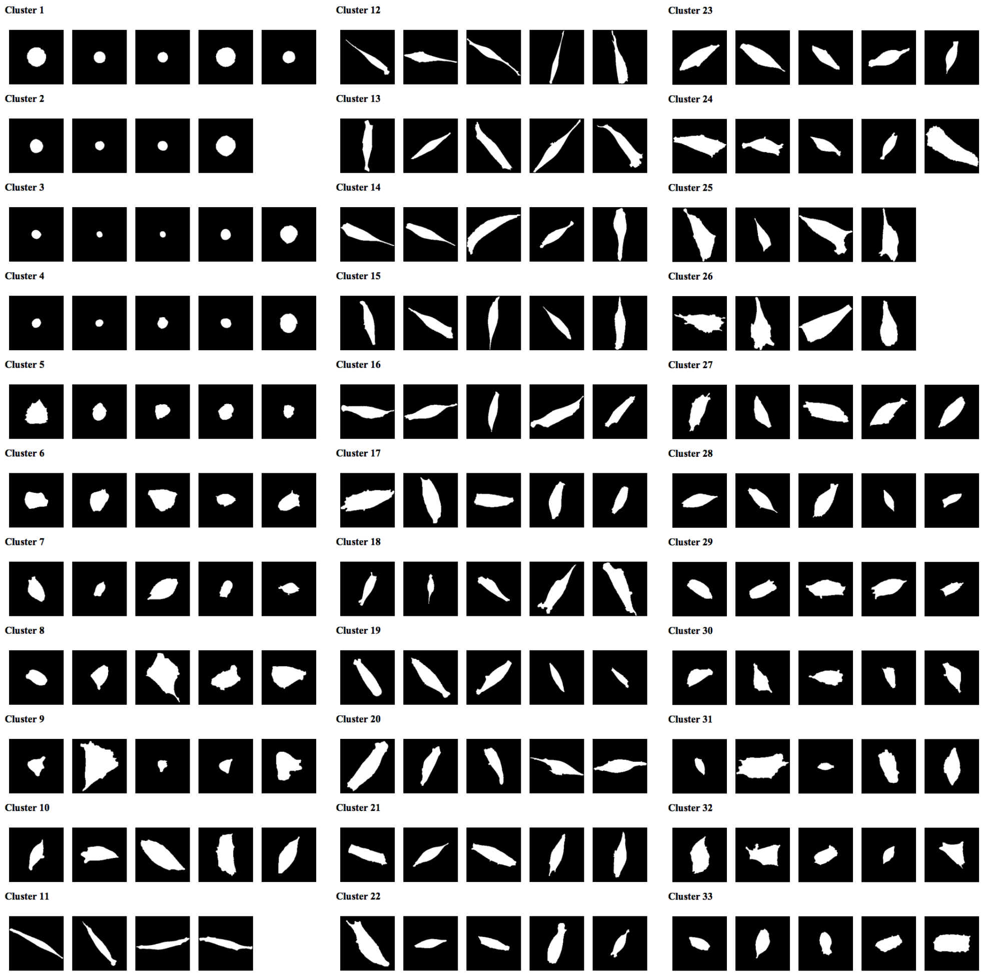


**Figure 4.** Clusters obtained by classifying cell shapes using Hu's invariant moments.

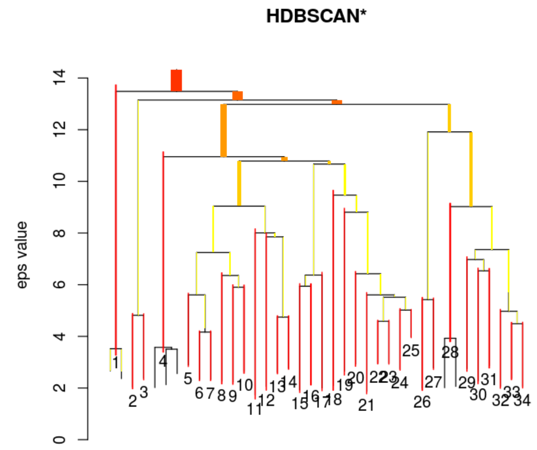


(a)

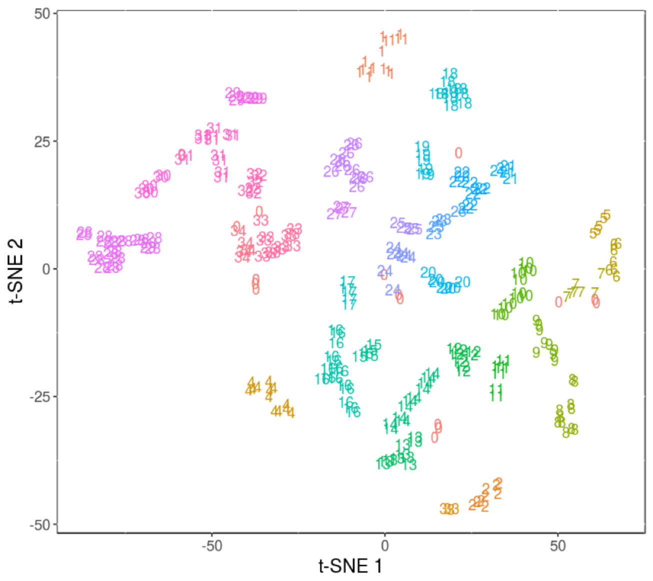


(b)

**Figure 5.** Clustering shape factors in a 2-component embedding
(a) Hierarchichal clustering of shape factors using HDBSCAN. (b) Cluster assignment. Cluster ID 0 indicates outliers.

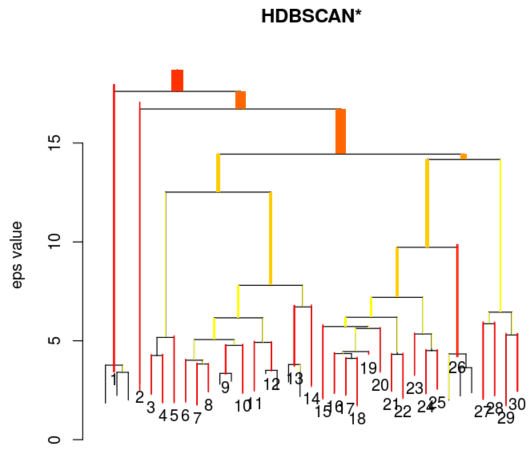


**(a)**

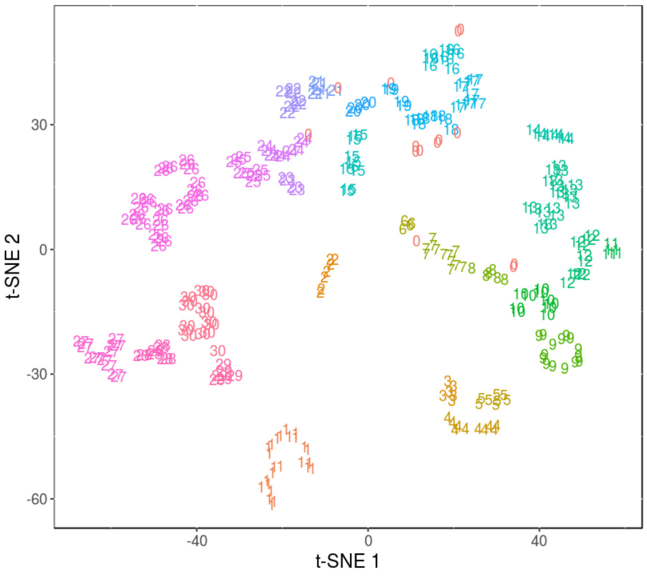


**(b)**

**Figure 6.** Clustering geometrical features in a 2-component embedding
(a) Hierarchichal clustering of geometrical features using HDBSCAN. (b) Cluster assignment. Cluster ID 0 indicates outliers.

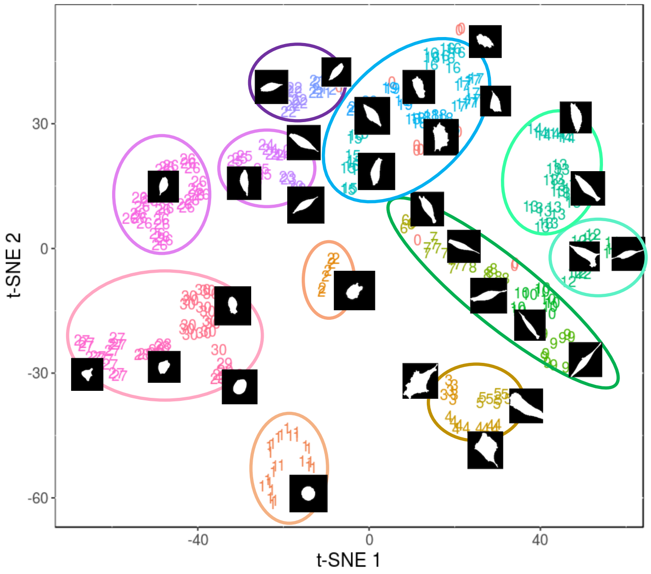


**Figure 7.** The 2D t-SNE plane, as in Fig. 6, but with closely related clusters (according to HDBSCAN) grouped, and with superimposed shapes of cells in each cluster.

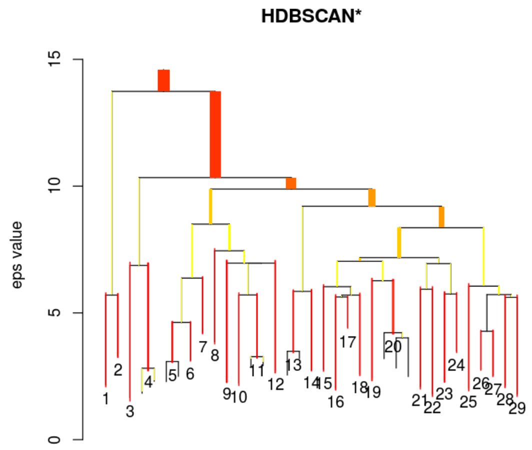


(a)

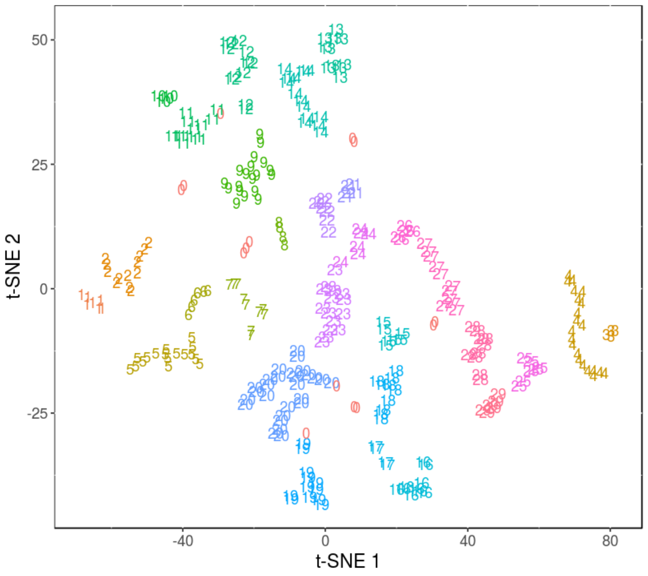


(b)

**Figure 8.** Clustering geometrical and boundary features in a 2-component embedding (a) Hierarchichal clustering of geometrical and boundary features using HDBSCAN. (b) Cluster assignment.

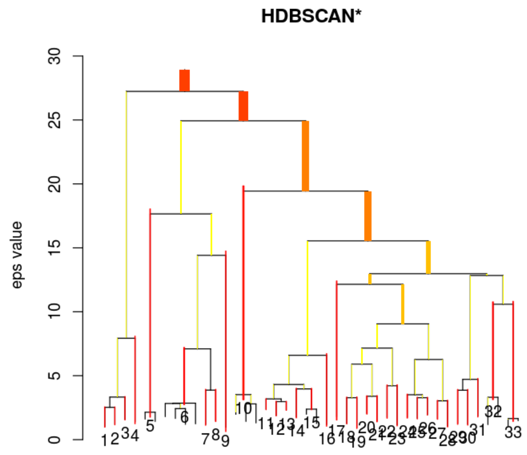


**(a)**

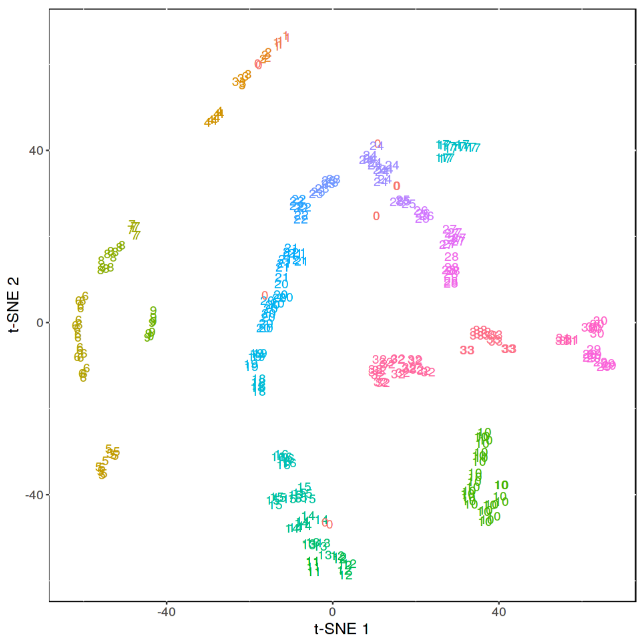


**(b)**

**Figure 9.** Clustering Hu's invariant moments in a 2-component embedding
(a) Hierarchichal clustering of geometrical and boundary features using HDBSCAN. (b) Cluster assignment.
